# A single genetic innovation at the origin of plant terrestrialization

**DOI:** 10.64898/2026.08.14.744897

**Authors:** Duchesse-Lacours Mbadinga Zamar, Philippe Ranocha, Mélanie Rich, Katharina Melkonian, Tatiana Vernié, Tifenn Pellen, Nicolas Vigneron, Jean Keller, Frédéric Domergue, Yves Martinez, Aurélie Le Ru, Christophe Dunand, Pierre-Marc Delaux

## Abstract

How the functional innovations that enabled plants to first colonize land 450 million years ago evolved remains puzzling. Here, we show that the gain of a single family of enzymes was pivotal in the evolution of two of these innovations, the cuticle that protect plants from dehydration and UV light, and the Arbuscular Mycorrhizal symbiosis promoting water and nutrient uptake. We show that functional phosphatase domain Glycerol-3-Phosphate Acyl Transferase (p-GPATs) evolved in the first land plants, concomitantly with the cuticle and symbiosis. Mutation of two p-GPATs from the liverwort *Marchantia paleacea* is sufficient to abolish both cuticle formation and AM symbiosis, leading to major developmental defects. We propose that the evolution of p-GPATs in land plants acted as a *two birds-one stone* innovation, diverting intracellular lipids to the extracellular space and providing a simple path to the evolution of two traits essential for the colonization of land.

## Introduction

The colonization of land by plants 450 million years ago was a critical event in Earth history.^1^ At the Earth-system level it led to a drastic decrease in atmospheric CO_2_.^1–3^ At the local level, terrestrial plants (Embryophytes), as primary carbon fixers, provided a resource for other organisms to thrive.^2,4,5^ For the ancestor of the Embryophytes, colonizing this new environment from freshwater meant overcoming new challenges, such as increased solar UV radiations, desiccation and nutrient deficiency. The first Embryophytes adapted to these obstacles through the evolution of new traits.^6^ For instance, the cuticle, a hydrophobic lipid and waxes layer at the plant surface, provided UV, pathogens and drought tolerance,^7–10^ while the mutualistic association formed between these first land plants and soil fungi, the arbuscular mycorrhizal (AM) symbiosis, improved nutrient and water uptake.^11–13^

Our understanding of the steps that led to the accumulation of all those innovations is hindered by the wide evolutionary gap between Embryophytes and their close algal relatives the zygnematophyceae and the scarcity of fossils of soft-tissue organisms. To fill this gap, we need to rely on what we can infer from the biology of the Most Recent Common Ancestor (MRCA) of the Embryophytes and try and reconstitute a likely chain of events leading to the cumulative adaptations of this organism.

Two traits recognized as major innovations leading to the terrestrialization are singular by their resemblance. The cuticle and AM symbiosis both rely on the production and export of lipids to the extracellular space, either the cell surface or the interface between the host and the symbiont cell.^10,14–17^ The cutin found in the angiosperm cuticle is a lipid polymer whose main precursors are Mono-Acyl-Glycerols (MAG). Production of these cutin monomers is mediated by plant-specific phosphatase domain-containing Glycerol-3-Phosphate Acyl Transferases (p-GPATs) that catalyze the addition of a fatty acid on the *sn-2* position of a Glycerol-3-Phosphate moiety, and the dephosphorylation of the *sn-3* position.^18–21^ On the other hand, AM symbiosis in Angiosperms relies on the transfer of large quantities of lipids from the host plants to the lipid auxotroph symbiotic fungi.^15,22,23^ Although the exact nature of the transferred lipids remains to be demonstrated, MAG are the primary candidates and REQUIRED FOR ARBUSCULAR MYCORRHIZATION 2 (RAM2,^24^), a p-GPAT specifically induced in the plant cells hosting the AM fungi, is essential for their biosynthesis.^13,15,16,23^ The analogy of mechanisms between those two traits led us to investigate their origin and conservation. Did those mechanisms co-occur in the MRCA of the Embryophytes or did one trait derive from the other in the evolution of angiosperms?

Using an evo-devo framework, including phylogenetics, trans-complementation assays and reverse genetics, we show here that p-GPATs belong to a family specific to land plants, and that their role in both cutin formation and AM symbiosis has been conserved across Embryophytes since their MRCA that lived on Earth 450 million years ago. The ancestral state being defined, we propose a ‘two birds - one stone’ model for the evolution of those two key terrestrialization innovations.

## Results and discussion

### p-GPATs evolved in the first Embryophytes

To precisely determine the evolutionary history of the p-GPAT family in the Viridiplantae lineage, we conducted a phylogenetic analysis using a database of 438 plant species, including 400 Embryophytes, covering Tracheophytes and Bryophytes which are the two main Embryophyte lineages, and 39 green algae encompassing all available genomes of streptophyte algae (Table S1). From the initial blast search, not a single sequence containing both an acyltransferase and a phosphatase domain were detected in any of the algal genomes, while all Embryophyte species harbored several homologs (Table S2). This confirms, with the largest available sampling, previous analyses proposing that p-GPATs are a specificity of the Embryophytes (Figure S1).^8,19,21,25^ The resulting phylogenetic tree resolved four main clades. The first clade encompasses sequences from all sampled lineages, including the green alga *Chlamydomonas reindardtii*, and the Angiosperm *Arabidopsis thaliana* GPAT9 (AtGPAT9). Proteins found in this clade are key players in the production of lipids for the primary metabolism.^26–28^ The other three clades are exclusively composed of p-GPATs from Embryophytes (Figure 1A), and include respectively AtGPAT1/2/3, AtGPAT4/6/8 and AtGPAT5/7 - *Medicago truncatula* RAM2 (MtRAM2). A well resolved clade of Bryophytes is found in the GPAT1/2/3 clade. Genes of this clade had been shown to be required for tapetum development or for the formation of cutin in the root cap.^29–31^ In addition, two bryophyte clades were identified at the base of the AtGPAT5/7-MtRAM2 and AtGPAT4/6/8 clades. They include the RAM2A, RAM2B and RAM2C paralogs from the liverworts *Marchantia polymorpha* and *Marchantia paleacea*.^13^ Our analyses could not resolve whether the duplication leading to these two bryophyte clades occurred before the diversification of the Embryophytes (*i.e.* before the split between Bryophytes and Tracheophytes) or after. The AtGPAT4/6/8 clade is generally known for its function in cutin biosynthesis in either leaves (GPAT4/8) or flowers (GPAT6).^10,32^ The AtGPAT5/7-MtRAM2 clade contain both the symbiosis-specific p-GPAT (RAM2), and p-GPATs with a degenerated phosphatase involved in suberin production (GPAT5/7).^19,24,33^ Altogether, these phylogenetic analyses demonstrate that p-GPATs were present in the MRCA of the Embryophytes but not in their close algal relatives; and that multiple rounds of gene duplication led to the expansion of this gene family across the Embryophytes.

**Figure 1.**
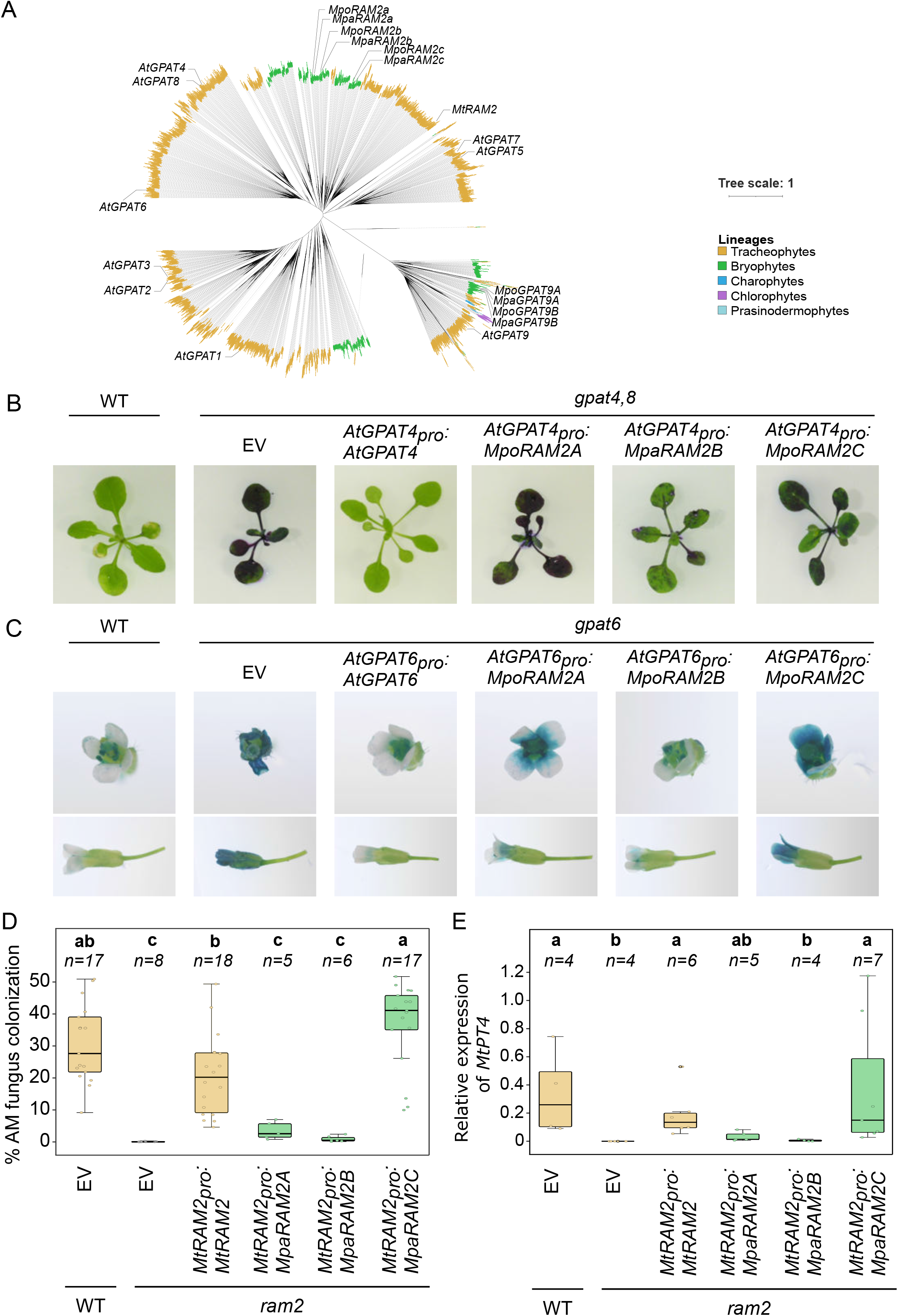
Conservation of p-GPAT across Viridiplantae. A. Unrooted phylogenetic tree of GPAT across green plants. Leaves are colored according to their lineage (orange: tracheophytes, green: bryophytes, blue: charophytes, pink: chlorophytes and cyan: prasinodermophytes). Genes from *Arabidopsis thaliana, Marchantia polymorpha, M. paleacea* and *RAM2* from *Medicago truncatula* are indicated. B. Dye penetration assays were carried out with toluidine blue on *Arabidopsis gpat4,8* plantlets complemented with various p-GPATs. Three independent lines were observed for each. See Figure S2 for a complete picture set. WT, wild type; EV, empty vector. C. Same as B for *Arabidopsis gpat6* flowers. See Figure S3 for a complete picture set. D. Arbuscular mycorrhizal symbiosis assays in *M. truncatula*. Wild-Type (WT) plants or *ram2* mutant roots transformed with an empty-vector (EV) control or with various p-GPATs. E. Relative expression of the symbiotic gene *PT4* in *M. truncatula* WT plants or *ram2* mutant transformed with an empty-vector (EV) control or with various p-GPATs, six weeks post inoculation with *R. irregularis*.

The literature suggests that p-GPAT biochemical functions are partially conserved across the different gene subclades in Angiosperms.^10,20,24,34,35^ To investigate if this holds true for all land plant p-GPATs, we tested the trans-complementation of mutants from two angiosperms, *A. thaliana* and *M. truncatula*, affected in cuticle formation and AM symbiosis respectively, with bryophyte p-GPATS. Using the endogenous genes as positive controls, we expressed the three *p-GPAT* (*RAM2A*, *RAM2B* and *RAM2C*) pro-orthologs of both the “cutin” and “AM Symbiosis” clades from *M. polymorpha* and/or *M. paleaceae* in *A. thaliana gpat6*, *gpat4/8* and *M. truncatula ram2* mutants under native promoters.

For all three mutants, at least one of the three Marchantia paralogs was able to rescue the developmental or symbiotic defects (Figure 1). *MpaRAM2B*, and to some extent *MpoRAM2B* and *MpoRAM2C*, rescued the rosette leaf impermeability caused by the cutin defect in the *A. thaliana gpat4/8* mutant (Figure 1B, Figure S2), while *MpoRAM2A* and *MpoRAM2B* complemented partially the petal permeability to toluidine blue of the *A. thaliana gpat6* mutant (Figure 1C, Figure S3). In *M. truncatula ram2* mutant, expression of *MpaRAM2C* was also able to rescue AM symbiosis colonization by the AM fungus *Rhizophagus irregularis* to a level similar to wild-type plants (Figure 1D). The expression of *MtPT4* (Figure 1E), a gene encoding for a symbiosis-specific phosphate transporter,^36^ was quantified by qRT-PCR and confirmed the functionality of the symbiosis in the *M. truncatula ram2* roots complemented with *MpaRAM2C* (Figure 1). By contrast, *MpaRAM2A* and *MpaRAM2B* did not rescue the symbiotic defects of the *M. truncatula ram2* mutant (Figure 1D-E)

Altogether, these data indicate that p-GPATs evolved in Embryophytes by duplication of an ancestral form of GPAT, followed by the integration of a phosphatase domain. The molecular properties of this new family of p-GPAT has been maintained for 450 million years. The ancestral GPAT is present in all eukaryote lineages, mediating the first step of the Kennedy pathway by esterifying the *sn-1* position of Glycerol-3-phosphate, leading to the biosynthesis of lysophosphatidic acid. By contrast, the functions of p-GPATs for Cutin biosynthesis and AM symbiosis require an active phosphate domain leading to the biosynthesis of Mono-Acyl-Glycerols.^18–21,24,34^ While the exact biological function of every p-GPAT genes is still to be determined, the published data strongly link genes of this family with either lipid transfer to a symbiont, or the biosynthesis of cutin and/or suberin monomers.^10,20,24,30–34^ Experiences in yeasts showed that Arabidopsis p-GPAT1/5 and 7 – that secondarily lost their phosphatase activity – but not p-GPAT2/3/4/6/8 are able to rescue a *gat1/gat2* mutant.^37^ This would suggest that the dephosphorylation of the sn2-glycerol-3-phosphate to MAG plays an important role in diverting the flux of lipids from the primary metabolism, a feature that would have evolved in the MRCA of the Embryophytes.

### The role of p-GPAT in cuticle formation is ancestral in land plants

If the evolution of p-GPATs in the first land plants indeed enabled the evolution of lipid export, the known biological function of p-GPATs in Angiosperms would be conserved in Bryophytes. Gametophores of a single *gpat2* mutant in the moss *Physcomitrium patens* displayed significant, yet very quantitative, cutin defect,^21^ supporting this hypothesis. Focusing on the three Marchantia p-GPATs from the Cutin/Suberin/AM symbiosis clades, we first assessed their expression pattern using promoter:GUS fusions in *M. polymorpha*. All three genes were expressed in the epidermis (Figure S4), with a stronger GUS staining at the margins of the air pores which are known to accumulate cutin in liverworts.^38^ In addition, staining was also observed in the lower epidermis (Figure S4). A similar expression pattern was observed for the three *RAM2* paralogs in *M. paleacea*,^13^ a pattern which is coherent with a putative function in cutin biosynthesis. To determine the function of p-GPATs in Marchantia, we generated both single and double mutants of the genes with the highest epidermal expression in *M. paleacea* using CRISPR/Cas9 (Figure S5). The single *Mparam2b* and *Mparam2c* mutants displayed a wild-type phenotype, although some *Mparam2c* alleles had a slightly delayed growth (Figure 2A, Figure S6). In contrast to the single mutants, the double *Mparam2b/ram2c* mutants displayed strong developmental phenotypes (Figure 2), with stunted growth, no lateral development and a lack of branching leading to a tube-like shape of the thalli (Figure 2A-B). This developmental defect is reminiscent of abnormal organ formation and organ fusion found in cutin defective *A. thaliana* mutants.^32,39–41^ Surface Electron Microscopy revealed that air pores were not formed on the surface of the *Mparam2b/ram2c* mutant alleles (Figure 2C, Figure S7). We conclude that *RAM2B* and *RAM2C* are partially genetically redundant and contribute to the development of *M. paleacea*.

**Figure 2.**
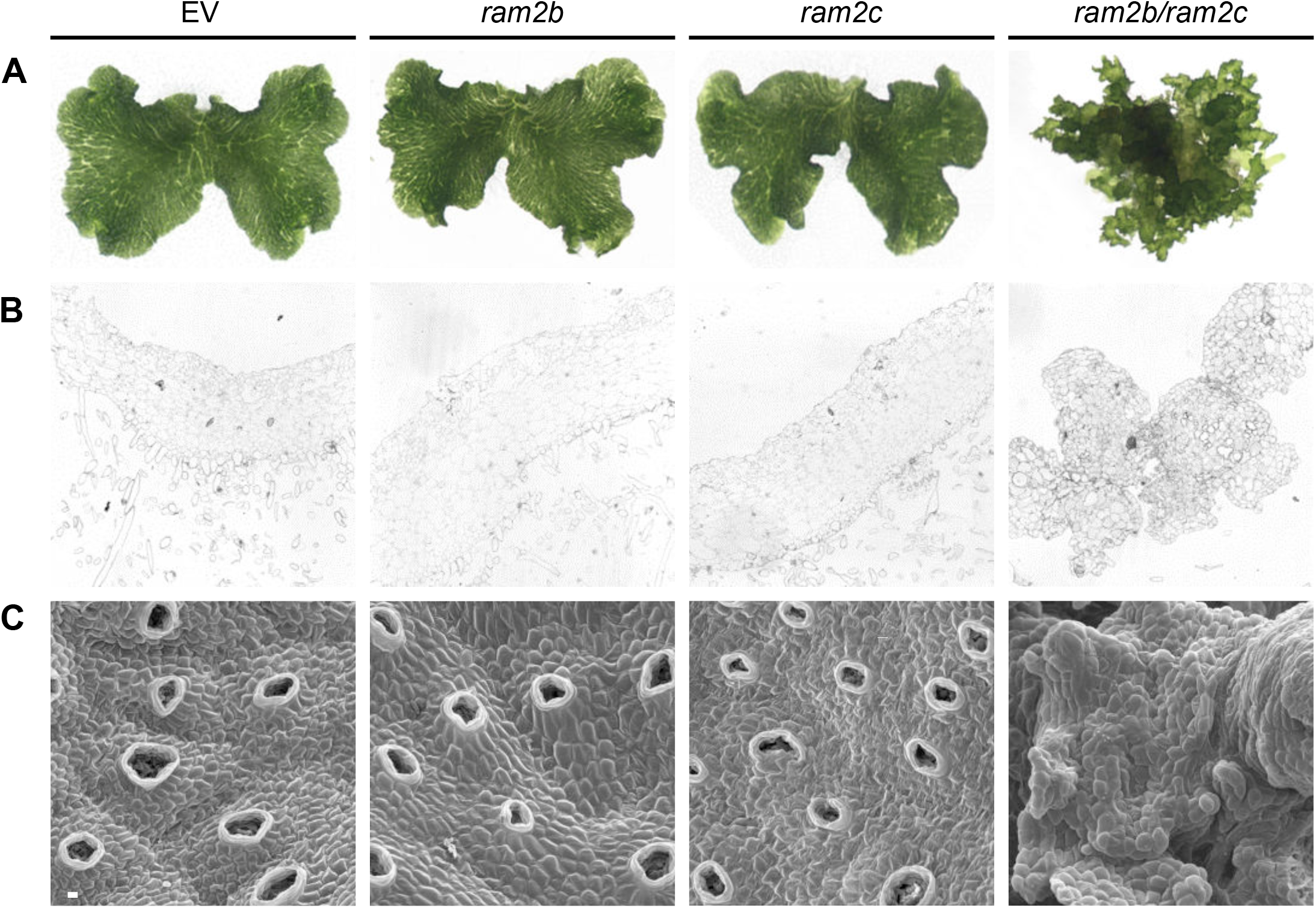
*RAM2B* and *RAM2C* contribute to *Marchantia paleacea* development. A. Overview of the wild-type (WT), single *Mparam2b* and *Mparam2c* mutants, and double *Mparam2b/ram2c* mutant morphology. Scale bar: 2 mm. B. Cross sections (1 µm) of the same lines embedded in Epon and stained with toluidine blue. Scale bar: 100 µm. C. Scanning electron micrographs of the dorsal face of the same lines. Scale bar: 35 µm. Three independent lines were observed for each. See Figure S7 for a complete picture set.

To directly observe the role of *RAM2B* and *RAMC* in cuticle biosynthesis, we visualized the cuticle in both *Mparam2b* and *Mparam2c* single mutants, and in the *Mparam2bram2c* double mutant using Transmission Electron Microscopy, and we quantified cutin monomers by GC-MS. The cuticle of both *Mparam2b* and *Mparam2c* mutants was indistinguishable from the wild-type plants (Figure 3A-B). Although a slight decrease in the abundance of some cutin monomers was observed for the *Mparam2c* mutant (Figure 3C), it did not result in observable cuticle defect (Figure 3A-B). In sharp contrast, in three independent *Mparam2b/ram2c* mutant lines, cuticle was not detectable, or only as a very thin layer (Figure 3A-B, Figure S8). Mirroring this defect, mass spectrometry analyses revealed a significant reduction (between 60% and 93% depending on the monomers) of cutin monomers in the three *Mparam2b/ram2c* double mutants (Figure 3C, Table S3). To further test the link between cuticle defects and the observed developmental phenotypes, we complemented one of the *Mparam2b/ram2c* double mutant alleles with *MpaRAM2c* under the control of its native promoter, or the epidermis-specific (Figure S9) *MpoSBG_pro_* promoter from *Marchantia polymorpha*.^42^ In both cases, developmental defects were rescued (Figure 3D), as well as the biosynthesis of cutin monomers (Figure 3C). This indicates that *RAM2B* and *RAM2C* are collectively essential for cutin biosynthesis in the epidermis, a function necessary for *M. paleacea* normal development.

**Figure 3.**
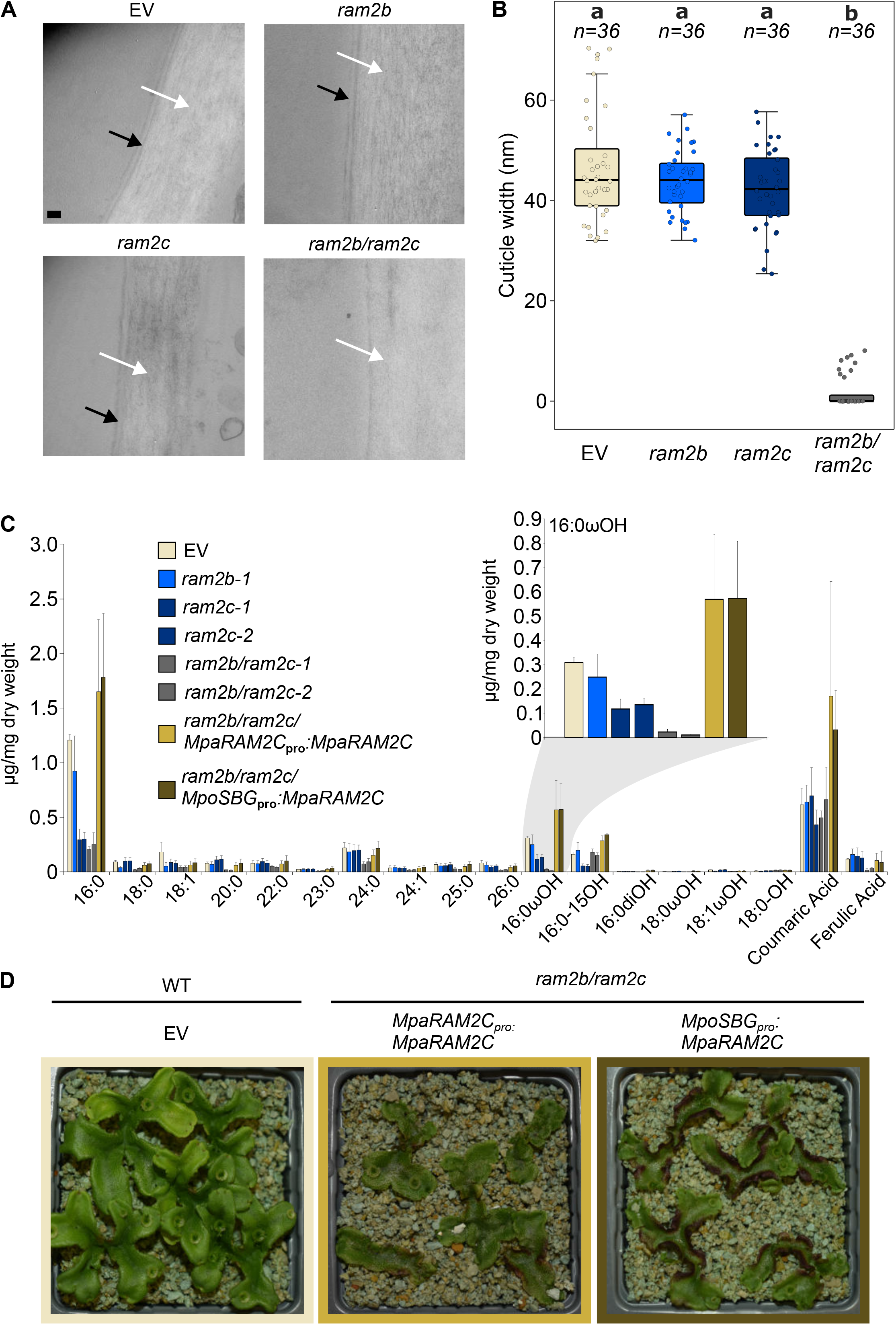
The p-GPATs *RAM2B* and *RAM2C* are essential for normal cuticle development in *Marchantia paleacea*. A. Epidermis of the wild-type *Marchantia paleacea*, single *Mparam2b* and *Mparam2c* mutants, and double *Mparam2b/ram2c* mutant as seen by transmission electron microscopy. Black arrow, cuticle; white arrow, cell wall. Scale bar: 100 nm. Three independent lines were observed for each. See Figure S8A for a complete picture set. B. Cuticle width (nm) of the same lines. See Figure S8B for a complete set of cuticle width measurement. C. Major cutin monomers in *M. paleacea* control, *Mpram2* mutants and complemented lines. See Table S3 for the complete dataset. D. Representative pictures of *M.paleacea* control plants transformed with an empty vector (left), and complementing lines in Mpa*ram2b/ram2c* mutant background, transformed with either Mpa*RAM2Cpro:MpaRAM2C* (middle) or Mpo*SBGpro:MpaRAM2C* (right).

The occurrence of p-GPATs specifically in Embryophytes, and their conserved function in cutin development, support the hypothesis that extracellular lipid polymers are a terrestrialization innovation.^8,21,25,42^ In an attempt to further test this hypothesis, we searched for the presence of a hydrophobic, lipid-based, polymer on the surface of four Zygnematophyceae using Nile-red staining and fluorescent microscopy (Figure S10). While Nile-red staining, indicative for the presence of neutral lipids, was observed inside cells of all four species, staining was never observed on the cell surface for any of the four algae, irrespective of the tested condition (Figure S10, see material and methods). We also analyzed these species for the presence of putative cutin monomers using hydrolysis followed by GC-MS analysis. In parallel, three liverwort species and one fern were used as diverse control representative of the Embryophytes. Mirroring the results obtained with *M. paleacea*, all liverworts and the fern displayed cutin monomers including C16:0-OH (Table S3,^42–44)^. By contrast, none of the typical hydroxylated cutin monomers were detected in any of the Zygnematophyceae (Table S3). Absence of a detectable cuticle layer in the algal clade sister to the Embryophytes which also lack p-GPATs reinforces the hypothesis that the evolution of the cuticle and p-GPATs are correlated.

Altogether, these data demonstrate that GPATs are essential for cutin biosynthesis in Bryophytes, as they are in Angiosperms, identifying p-GPAT-mediated cutin formation as an ancestral feature of land plants.

### The role of GPAT in AM symbiosis is ancestral in land plants

The transfer of lipids from host plants to arbuscular mycorrhizal fungi has been identified as one of the essential features of AM symbiosis that was present in the MRCA of the Embryophytes.^13,15,16,23,45^ As indicated above, in Angiosperms such as *M. truncatula* or rice,^24,34,46^ this symbiotic lipid transfer requires the p-GPAT RAM2. However, the symbiotic function of RAM2 beyond angiosperms has never been investigated, leaving open the question of its origin. In *M. paleacea*, RNAseq and promoter:GUS studies indicated that *MpaRAM2B* and *MpaRAM2C* are induced in arbuscule-containing cells, where lipid transfer occurs.^13^ To determine whether the function of RAM2 in symbiosis is conserved across land plants, we inoculated *M. paleaecea Mparam2b, Mparam2c* and an empty vector control with spores of the symbiotic fungus *Rhizophagus irregularis*. After five weeks, 85% of the wild-type plants showed the typical pigmentation induced by an established symbiosis (Figure S11). Colonization was confirmed by staining of the AM fungus in sections of the control thalli (Figure S11B). Quantification and microscopy analyses of the independent *Mparam2b* mutant alleles did not identify symbiotic defects (Figure S11). While in some experiments one of the *Mparam2c* alleles displayed reduced colonization, this phenotype was not robust across three independent replicates and alleles (Figure S11). Given the massive developmental phenotype of the double *ram2b/ram2c* mutant, we did not include it in the AM symbiosis phenotyping analysis. Instead, we took advantage of the *Mparam2b/ram2c* double mutant complemented with the epidermis-specific *MpoSBG_pro_:RAM2C or MpaRAM2c_pro_:MpaRAM2C* constructs. Complementation of the *Mparam2b/ram2c* double mutant with *MpaRAM2c_pro_:MpaRAM2C*, driving *MpaRAM2C* expression in both the epidermis and the area colonized by *R. irregularis*, resulted in slightly reduced colonization rates compared to an empty vector control when quantified as number of thalli per genotype showing the AM symbiosis-induced pigment (Figure 4A). Colonization rates were similar in the lines complemented with *MpoSBG_pro_:RAM2C* (Figure 4A), which were rescued for their developmental defects to different extent, ranging from some rhizoids being produced at the base of tube-like structures, to plants with fully-restored thalloid development. Sections of *the Mparam2b/ram2c* double mutant complemented with *MpaRAM2c_pro_:MpaRAM2C* stained with WGA-Alexafluor revealed the occurrence of normal fungal structures, including intracellular hypheae and fully-formed arbuscules, indicative of a functional symbiosis (Figure 4A-B). In sharp contrast, confocal imaging of the colonized area following WGA-Alexafluro staining in the *MpoSBG_pro_:RAM2c* complemented-*Mparam2bram2c* double mutants revealed the total absence of arbuscules in most lines, and eventually some collapsed and unbranched arbuscules in others (Figure 4), which is reminiscent of the symbiotic phenotypes observed for the *M. paleacea wri* mutant impaired in the activation of the symbiotic lipid-transfer pathway,^13^ or the *ram2* mutants in angiosperms.^23,24,34,46^

**Figure 4.**
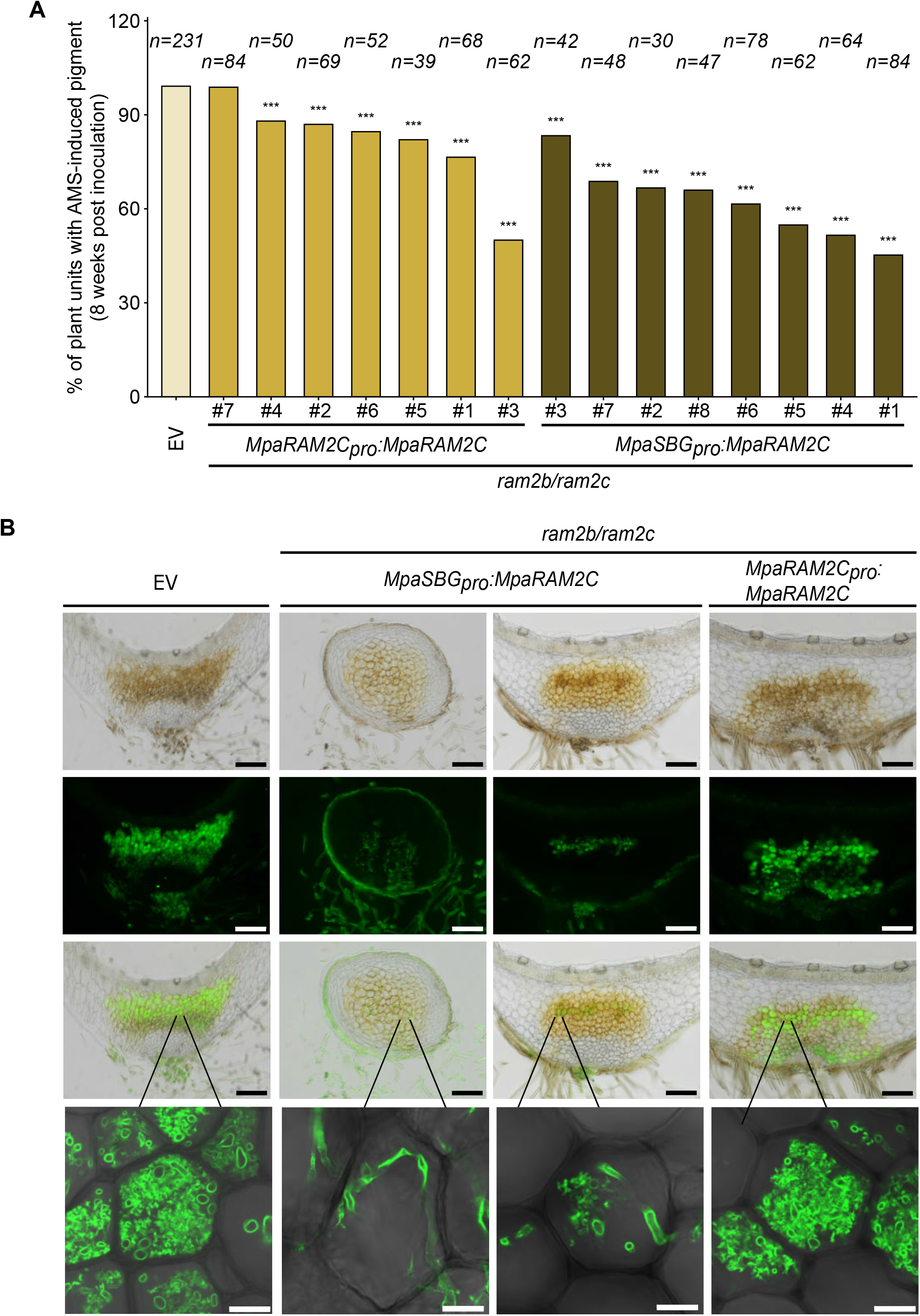
p-GPATs are essential for functional Arbuscular Mycorrhizal symbiosis in *Marchantia paleacea*. A. Scoring of plant units with AMS-induced pigment at 8 weeks post inoculation with *R. irregularis* in *M. paleacea* control plants (left), and complementing lines in Mpa*ram2b/ram2c* mutant background, transformed with either Mpa*RAM2Cpro:MpaRAM2C* (middle) or Mpo*SBGpro:MpaRAM2C* (right) at eight weeks post inoculation with *R. irregularis*. Combined results of 3 independent experiments are shown. B. Representative images of *M. paleacea* control plants transformed with an empty vector (left) and complementing lines in Mpa*ram2b/ram2c* mutant background, transformed with either Mpo*SBGpro:MpaRAM2C* (middle) or Mpa*RAM2Cpro:MpaRAM2C* (right) inoculated with *R. irregularis* at 8 weeks post inoculation. From top to bottom: brightfield, green channel (WGA-Alexafluor 488 staining of fungal structures), merge, magnification. Scale bars are 250 μm or 20 μm for magnified images.

Altogether, these experiments demonstrate that the clade of p-GPATs involved in cutin formation in Bryophytes is also essential for functional AM symbiosis. Hence, integrating this discovery and the knowledge from angiosperms, the most parsimonious hypothesis is that p-GPATs were already involved in both processes in the MRCA of the Embryophytes.

### Proposing a model for the evolution of lipid-export traits during terrestrialization

Although Bryophytes and vascular plants are very different in their morphology, they share a wide range of the mechanisms associated with their adaptations to the terrestrial environment. This includes for instance the developmental control of different tip-growing epidermal cell types,^47^ the formation of the cuticle ^8,10,21,32,42,48^ or symbiotic signalling.^13,49–51^ This inferred complexity of the MRCA of land plants suggests a very wide window of time during which the first land plants evolved to gain those traits. Neither an extant species nor fossil specimens have yet been discovered that could provide information on intermediary stages leading to this complex MRCA. Yet, likely sequences of events can be theorized.

A consensus of ancestral state reconstructions is that the MRCA of land plants and streptophyte algae was a filamentous multicellular organism,^52,53^ but all embryophytes show a 3D “tissue” structure and dissections of the underlying mechanisms confirm this transition occurred before land plant diversification.^54–57^ We can hypothesize that the transition from a filament to a 3D shape arose early in the adaptation of plants to aerial conditions as this kind of structure minimize the surface/volume ratio and the potential water loss (State 1, Figure 5).

**Figure 5.**
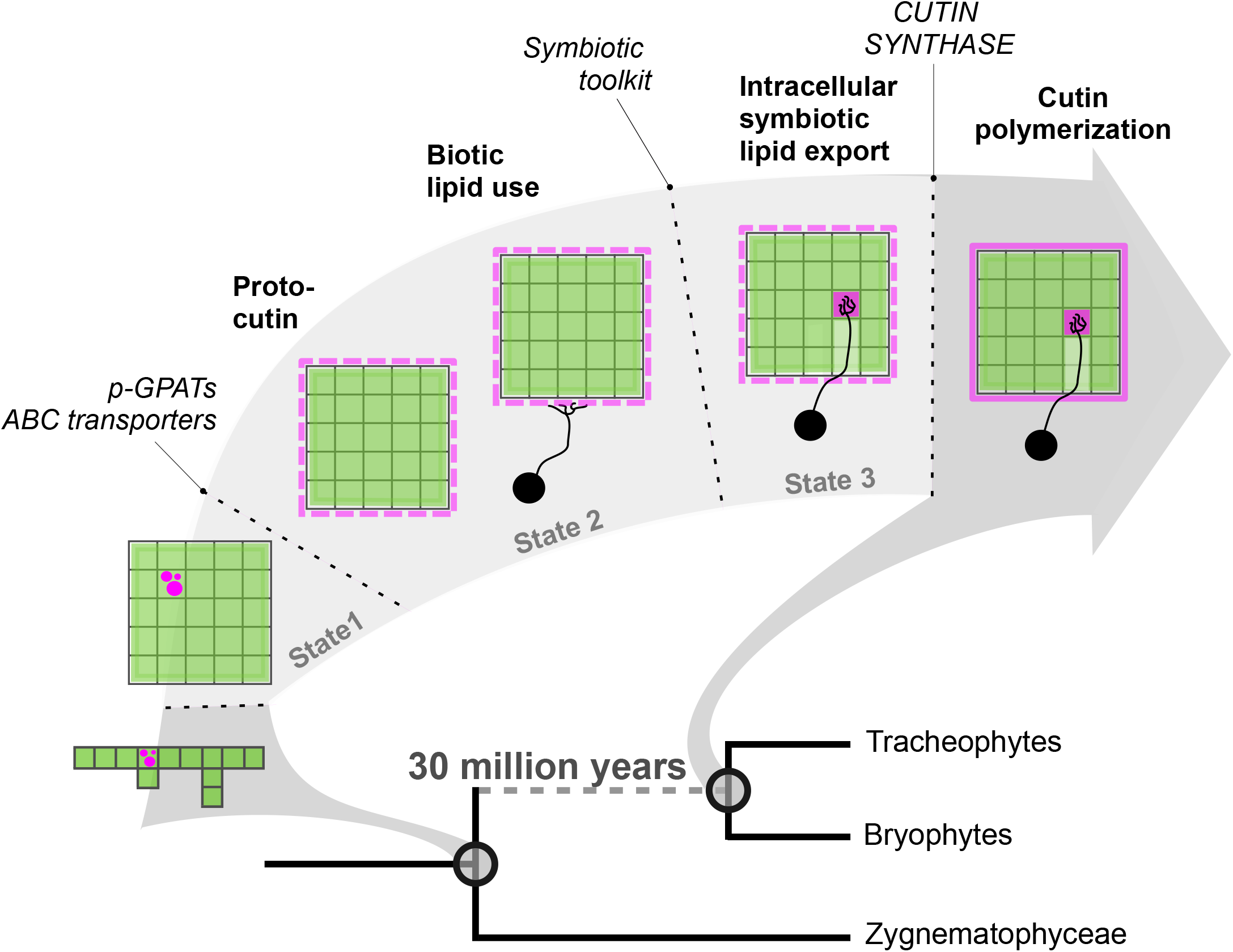
A stepwise model for the evolution of cutin and Arbuscular Mycorrhiza Symbiosis.

In the algal cousins of land plants, lipid accumulation, in the form of triacylglycerols (TAG) is a common stress response, especially in the case of nutrient deficiency.^58,59^ The conditions faced by first the land plants are likely to have triggered such a response. We propose that a decisive step came with the addition of a mechanism of lipid export, combining p-GPATS synthesizing a novel type of lipid intermediate and plasma membrane transporters. These excreted lipids would have readily provided direct benefits in terms of UV and desiccation tolerance. The basis of the extracellular transfer of these monomers remains elusive, even in extant land plants, although ABC transporters are likely candidates.^14,60^ ABC transporters (STR and STR2) are also thought to be involved in the symbiotic transfer of lipid,^61^ suggesting that the export mechanism might have evolved only once, probably from gene families already diversified in streptophyte algae.^62^ We propose the biosynthesis and export of sn2-MAGs as the second intermediate state toward the evolution of the MRCA of land plants (Figure 5).

Excreted lipids also represent an attractive carbon source for terrestrial microbes. The most common road for a microorganism to become a mutualistic partner is to start from being saprotrophs or pathogens.^63,64^ It is likely that the fungi the first plants encountered on land preyed on decaying poorly adapted organisms. But plants producing extracellular lipids might have been a gateway for the fungi to profit from the photosynthetic abilities of plants in a non-destructive way. Providing those plants with mineral nutrient and water maximized this carbon output. In the intermediate state 3, the plant and the fungal partners would have been engaged in a mutually beneficial interaction sharing goods in their direct environment (State 3, Figure 5).

Refinement of the cuticle, , including the evolution of the CUTIN SYNTHASE,^65^ led to a polymerized hydrophobic layer improving resistance to UV and desiccation. However, the shift in availability of the excreted lipids probably affected the evolving mutualism. We propose that the evolution of a polymerized cuticle was contingent upon the prior gain of the ability to spatially restrict and ultimately internalize fungal hypheae (Intermediate state 4), resolving this bottleneck while providing opportunity for proper symbiont control/checkpoint and creation of the very large surface of exchange that is the periarbuscular space.^66^ Finally, the consecutive loss of the fatty acid synthase in Glomeromycota effectively stuck AM fungi in the mutalist end of the symbiosis spectrum.^22,67^

## Conclusion

Experimental evolution and theoretical work propose that trait evolution relies on three successive steps: potentiation, actualization and refinement.^68,69^ From inferences based on the fossil record and extant species it can be proposed that the traits deemed necessary for plant terrestrialization, such as cuticle and AM symbiosis, were already in place - refined - and regulated by complex pathways in the MRCA of the Embryophytes.^7,8,11,13,25,70–72^ Our reverse genetic analysis of the p-GPATs in *M. paleacea* embedded in an evo-devo framework demonstrates that both cutin formation and lipid transfer during AM symbiosis are ancestral in Embryophytes. More surprisingly, we demonstrate that they are homologous traits. We propose an evolutionary model in which the fusion of a phosphatase domain and the pre-existing Acyl-Transferase domain acted as a potentiation event, providing the key enzyme to redirect the central lipid metabolism to extracellular routes, and providing the building blocks for two of the terrestrialization innovations to subsequently evolve. Beyond the plant colonization of land, we conclude that trait evolution, although depending on a multitude of refinements before reaching a fixed state, may arise from a combination of few large-effect molecular changes.

## Acknowledgements

The authors thank the genotoul bioinformatics platform Toulouse Occitanie (Bioinfo Genotoul, https://doi.org/10.15454/1.5572369328961167E12) for providing computing resources, and the Bordeaux-Metabolome Facility, which is supported by MetaboHUB (ANR-11-INBS-0010), where cutin analyses were performed. We are grateful to Bruno Payre of the Centre de Microscopie Electronique Appliquée à la Biologie (Faculté de Médecine Rangueil, Toulouse, France) for his assistance. This project has received funding from the European Research Council (ERC) under the European Union’s Horizon 2020 research and innovation program (grant agreement No 101001675 - ORIGINS) to P-M.D. This work was supported by the “Laboratoires d’Excellence (LABEX)” TULIP (ANR-10-LABX-41)” and by the “École Universitaire de Recherche (EUR)” TULIP-GS (ANR-18-EURE-0019), by a PhD fellowship from the University of Toulouse to N.V., and a Postdoc fellowship from the German Research Foundation (DFG Walter Benjamin fellowship project number 536856410) to K.M.

## Materials and Methods

### GPAT orthologs data mining and Phylogeny

GPAT orthologs were retrieved from a database containing 438 plant species covering the diversity of Viridiplantae using the reference proteins of *Arabidopsis thaliana* nine GPATs and the BLASTp+ v2.15.0 with a e-value threshold of 1e-05 and a maximum of 3000 target sequences reported for each query.^73^ Protein sequences were then aligned using MAFFT v7.505 with the ‘—auto’ option and a maximum of 30 iterations.^74^ Alignment was then subsquently trimmed using trimAl v1.4.1 to remove positions containing more than 60% of gaps.^75^ Phylogenetic reconstruction was performed using FastTree v2.1.11 with default parameters and the ‘-gamma’ option enabled.^76^ Tree was visualized and annotated within the iTOL v7 platform.^77^ Functional domains within each GPAT orthologs were identified using HMMSCAN from the HMMER v3.4 package with the e-value thresholds for both model and domain reporting set at 1e-04.^78^

### Confocal microscopy and histochemical analyses

Cutin was visualized on one-week-streptophyte algae cells and on fresh 100μm size transversal sections of one-month-old Marchantia thalli. Samples were first stained with a primary cell wall-labeling solution containing 0,01% Calcofluor White in acetone for 1min (streptophyte algae) or 2min (Marchantia). Then, samples were rinsed with ultra-pure water and incubated in the lipid tagging solution containing 0,001mg.ml^-1^ Nile red in acetone for 2min. Following the incubation of samples in the Nile-red solution, the marked samples were immediately mounted between slide and cover glass for microscopic imaging. Images were captured with a LEICA SP8 confocal laser scanning microscope. The Calcofluor excitation wavelength used was 405nm and the fluorescence signals for the emission were detected from 415 to 485nm. The chlorophyll auto-fluorescence was observed with an excitation wavelength of 552nm, and the emission of fluorescence signals was detected from 630 to 670nm. The Nile-red excitation used was 488nm and emission of fluorescence signals was from 550 to 640nm.

Histochemical GUS analyses were performed on five independent transgenic lines for each promoter:GUS fusion construct. For *Marchantia polymorpha*, approximately, fifteen freshly detached gemmae from each transgenic line were transferred to soil and grown under the conditions described below for Marchantia culture. One-month-old thalli were carefully rinsed with ultra-pure water to remove residual plant substrate. For *Marchantia paleacea*, 2-weeks old transgenic plants were mock-inoculated or inoculated with *Rhizophagus irregularis* and grown side by side as described below for another seven weeks. The substrate was carefully removed and GUS expression in presence versus absence of the symbiont was assessed subsequently. GUS staining was performed by vacuum infiltrating thalli for 5 or 20 min in GUS staining buffer, followed by overnight incubation at 37°C. The staining buffer contained 0,5M EDTA, 50mM Potassium buffer, 100mM ferrocyanide potassium and 100mM ferricyanide potassium, 1% triton and, 2,5μg.ml^-1^ X-Gluc. After the overnight incubation, thalli were rinsed with ultra-pure water, embedded in 7% agarose, and transversely sectioned in 100μm slices using a LEICA VT1000 vibratome. Sections were subsequently incubated in 70% ethanol to remove chlorophyll and stored in the same solution until imagery. Images were acquired using a ZEISS Axio zoom V16 microscope or a Nikon Ti Eclipse inverted microscope equipped with DS Ri2 camera and motorized XY stage. Large images of global sections were generated using the NISAR 4.3 scan large image module allowing multifield acquisition and images stitching. These images were acquired with 10×/0.3 dry objective (0.73 pixel size) in brightfield and in fluorescence for WGA-Alexa 488 staining using a GFP band pass filter set (ex: 472/30 nm, em:520/35 nm).

### Transmission Electron Microscopy

Unless stated otherwise, all products were purchased from Electron Microscopy Sciences (EMS). Small sections (*ca*. 2 x 3 mm, midrib region) of fresh *Marchantia paleacea* thalli of approximately the same size were fixed for 1.5 hours at room temperature in a solution of 2% glutaraldehyde (EMS #16220) in 0.1M trihydrate sodium cacodylate buffer (EMS #12300) pH 7.2. Following fixation, the samples were washed 3 times (for 15 minutes each) with 0.1M trihydrate sodium cacodylate buffer pH 7.2, post-fixed with 1% OsO4 (EMS #19150) in cacodylate buffer for 2 hours at room temperature, and washed 3 times as previously.

Samples were dehydrated through an ethanol series (25% for 1h at room temperature, 50% overnight at 4°C, 50% for 30 minutes twice at room temperature, 70% overnight at 4°C, 90% for 1h at room temperature, 100% for 1h three times at room temperature). Uranyl acetate (1%, EMS #22400) was added to the first 50% ethanol step. After dehydration, samples were incubated twice (1h each) in propylene oxide (EMS #20410), and were gradually embedded in Epon812 resin (EMS #14120) mixed with propylene oxide at 1:2, 1:1, 2:1 ratio, each for 2h at room temperature and overnight at 4°C. Finally, samples were incubated twice in 100% Epon812 resin for 2h at room temperature and overnight at 4°C. Epon impregnated samples were cast in flat embedding molds (EMS #70903) and polymerization was carried out for 48 hours at 60°C (Reichert KT 100 oven).

Ultra-thin sections (80 nm) were cut from the embedded samples using a DiAtome diamond knife microtome (Reichert-Jung Ultracut E), and placed on formvar coated 100-mesh nickel grids (EMS #215-412-8400) for staining with Reynolds lead citrate (Delta Microscopies #11300) for 2 minutes under CO_2_-depleted atmosphere. Transmission electron microscopy images were captured at an accelerating voltage of 80 kV using a CCD camera on a Hitachi HT 7700 microscope. Cuticle width was measured with ImageJ (National Institutes of Health, https://imagej.nih.gov/ij/).

### Surface Electron Microscopy

Fresh *Marchantia paleacea* thalli were fixated and dehydrated through an ethanol series (up to 100%) as described above, except for the presence of uranyl acetate at the 50% ethanol step. Samples were dried in an EM CPD300 critical-point dryer (Leica Microsystems) using liquid CO_2_ as transitional fluid, mounted on aluminum stubs over double-sided carbon tape, sputter-coated (EM MED020, Leica Microsystems) with a 5 nm-thick platinum layer, and observed with a scanning electron microscope (FEI Quanta 250 FEG) at an accelerating voltage of 10 kV.

### Plants and growth conditions

Streptophyte algae species were obtained from CCAP (culture collection of algae & protozoa, www.ccap.ac.uk) and cultured under sterile conditions in soil extract-free liquid Woods Hole medium adjusted to pH 7.2, under an 8h light/16h dark photoperiod at 22°C.

*Marchantia polymorpha* was grown on soil by sowing gemmae under the same environmental conditions as those used for streptophyte algae.

*Marchantia paleacea* plants were grown on zeolite substrate (50% fraction 1.0 to 2.5 mm, 50% fraction 0.5 to 1.0-mm, Symbiom) in a walk-in growth chamber at 20 °C under 16 h light/8 h dark cycle.

For in vitro culture, gemmae of both Marchantia species were surface-sterilized in a 0.25% bleach solution for 30 seconds, followed by two or more rinses with sterile ultra-pure water to remove excess bleach. Sterile gemmae were then cultured on solid ½ Gamborg media (pH 5.6) 1.4% agar, with (*M. polymorpha*) or without (*M. paleacea*) 1% sucrose.

*Arabidopsis thaliana* accessions Columbia-0 (Col-0), *gpat4* x *gpat8* (SALK_106893 x SALK_095122, Li *et al*., 2007) and *gpat6-1* (SALK_136675, Li-Beisson *et al.*, 2009) were obtained from the Nottingham Arabidopsis Stock Centre (NASC). Homozygous mutant lines were identified by PCR using the gene-specific primers listed in Table S5.

*Arabidopsis* plants were routinely grown in Jiffy-7 (http://www.jiffygroup.com/) peat pellets (8h light, 200 µmol photons m^-2^ sec^-1^, 22°C, 67% relative humidity). For in vitro experiments, seeds were surface-sterilized and sown on agar-solidified (Euromedex #1330) half-MS medium (Sigma-Aldrich #M9274), in a culture room with an 8-h photoperiod (150 µmol photons m^-2^ sec^-1^) at 22°C.

*Medicago truncatula* A17 (wild-type) and *ram2* mutant were provided by the John Innes Center.^24^

### Genetic transformation

*Marchantia paleacea* transformation was conducted as previously described, using Hygromycin selection.^13^ For the complementation of the mutant line *ram2-2/ram2-3*, in absence of gemmae, one selected thallus was propagated by cutting it in pieces of ∼2mm in two rounds of cutting until sufficient amount of material was available. Selection of the retransformation was performed on 0.5µM Chlorosulfuron.^79^

For *Arabidopsis thaliana* transformation, constructs were introduced into *Agrobacterium tumefaciens* strain GV3101 (pMP90) for subsequent floral-dip transformation of *Arabidopsis*,^80^ albeit with Silwet L-77 (Lehle Seeds #VIS-01) concentration reduced to 0.002% to circumvent its apparent toxicity to some *gpat* mutant lines. Transformed seeds were selected on half-MS medium containing Hygromycin (12mg/l, Sigma-Aldrich #H0654). Three independently transformed homozygous lines were identified for each construct after several rounds of self-pollination.

Transformations of *Medicago truncatula* were performed following the published protocol^81^. After transformation, seedlings were transferred on Fahraeus medium and grown for 3 weeks at 23°C 16/8 photoperiod at 80–100 mmol m-2 s-1 provided by fluorescent bulbs. Transgenic roots were screened with an epi-fluoresence microscope. Seedlings with transgenic roots were selected based on DSred expression.

### Genetic constructs

To generate GUS reporter lines, approximately 2kb of the Marchantia *RAM2* promoter regions, a 35S terminator and the GUS coding DNA sequences were synthesized and inserted into a pL0M plasmid to generate level 0 modules. The modules were subsequently assembled into binary vectors for plant transformation using the Golden Gate modular cloning method. The pL1V-R2-47811 and pL2V-50507 backbones were used to assemble level 0 modules and level 1 parts, respectively. To test the ability of Marchantia genes to complement the symbiotic or cuticle phenotypes of Medicago and Arabidopsis mutants, a 2kb fragment of the native promoters (*MtRAM2*, *AtGPAT4* and *AtGPAT6*) from the relevant angiosperm species and the full-lenght CDSs from both angiosperms (*MtRAM2*, *AtGPAT4* and *AtGPAT6*) and bryophytes (*Mpa/MpoRAM2A*, *RAM2B* and *RAM2C*) were first synthesized and cloned as level 0 modules. The final genetic constructs were generated by assembling the promoters and CDS sequences into a binary vector for plant transformation following the same approach described above for GUS lines. Nucleotide sequences of genetic constructs generated at levels 1 and 2 are provided in Table S5.

*Marchantia paleacea* mutants of the *RAM2B* and *RAM2C* genes were generated using CRISPR/Cas9. Two guides targeting the *RAM2C* genes were used in a single construct and 3 independent alleles presenting different editions were selected (Table S5, Figure S5). To generate the double *ram2b/ram2c* mutant, constructs with 1 guide targeting each gene were used. The *ram2b* mutants were selected from the pool of transformed lines carrying the *RAM2B/RAM2C* guides but presenting with a wild type phenotype.

For the *ram2b/ram2c* complementation, *ram2b/ram2c-1* was transformed with the MpaRAM2C cDNA sequence under the native MpaRAM2C promoter (2kb, ^13^) or the promoter of the *M. paleacea* ortholog of MpoSBG9, a gene expressed in the upper and lower epidermis and involved in the regulation of cutin biosynthesis (2kb ^42^).

### Biochemical analysis

The biochemical diversity of cutin across streptophyte algae and Embryophytes was characterized using 0,5 to 2g of non-ground (3- to 4-weeks-old streptophyte algae) or pre-cut (1-month-old embryophytes) fresh plant samples. For cutin analysis of wild-type and mutant of *Marchantia paleacea* plants, 150 to 300mg of fresh tissues was used. Isolation of cutin extracts, depolymerisation, derivatization by sylilation, and GC‒MS identification / quantification were performed as described in^82^. Cutin monomers content of wild-type and mutant of *Marchantia paleacea* plants was expressed in µg/mg DR (dry residue) since surfaces could not be accurately measured. While for the biochemical diversity of cutin across streptophyte algae and Embryophytes data are expressed in µg/mg of fresh material as the algal dry mass was negligible after the drying step preceding the extensive delipidation procedure.

### Leaf and flower permeability tests

For epidermal permeability, *gpat4* x *gpat8* 3-week-old plantlets and *gpat6-1* flowers were incubated in 0.05% toluidine Blue O (Electron Microscopy Sciences #26074-15) with 0.01% Tween 20 (Sigma-Aldrich #P1379) for 10 minutes, and then abundantly rinsed in water before observation with a Zeiss Axio Zoom V16.

### AMS phenotyping

Five *M. paleacea* plants per pot were grown in a walk-in growth chamber at 20 °C under 16 h light/8 h dark cycle for 2 weeks. Each pot was then mock-inoculated or inoculated with approximately 1,000 sterile spores (250 spores per plant) of *Rhizophagus irregularis* DAOM 197198 provided by Agronutrition (Labège, France, https://www.agronutrition.com/en/contact-us/) and co-cultivated for another 8 weeks. All plants were watered once a week with Long-Ashton solution containing 15 μM of phosphate. At 8 weeks post inoculation, the thalli were harvested and cleared in pure ethanol to remove chlorophyll. Cleared *M. paleacea* thalli were scanned and scored as previously described using the presence of an AM symbiosis-specific black/purple pigment.^49^ To confirm presence or lack of fungal structures, the thalli were subjected to microscopic imaging. Thalli were embedded in 7% agarose and 100 µm transversal sections were prepared using a Leica vt1000s vibratome. The sections were incubated overnight in 12% KOH and washed 3 times with water. To visualize arbuscules, the sections were incubated in 1μg/ml WGA-AlexaFluor 488 (Invitrogen, France) in PBS buffer overnight at 4 °C in the dark. Sections were imaged using a Nikon Ti Eclipse inverted microscope equipped with DS Ri2 camera and motorized XY stage. Images were generated using the NISAR 4.3 scan large image module that allows multifield acquisition and image stitching. Images were acquired with 10×/0.3 dry objective (0.73 pixel size) in brightfield and in fluorescence for WGA-Alexa 488 staining using a GFP band pass filter set (ex: 472/30 nm, em:520/35 nm). In addition, close-up images were acquired using a Leica SP8 TCSPC confocal microscope and LAS X software with a 25× water immersion objective (Fluotar VISIR 25×/0.95 WATER) at zoom 5× (0.182 μm pixel size). WGA-Alexa 488 staining of fungal structures was excited with the 488 nm laser line and fluorescence was recovered between 500 nm and 550 nm. Brightfield images were also acquired and merged. Images were processed with ImageJ.

For *Medicago truncatula*, plants were grown in pots containing a mix of zeolite 1.0-2.5 mm (Symbiom) and sands (v/v), and inoculated with 300 spores of the *Rhizophagus irregularis* DAOM 197198 strain (Agronutrition). Plants were watered weekly with 0.5X Long Ashton medium containing 7.5 μM NaH2PO4. Roots were harvested four weeks after *R. irregularis* inoculation and washed three time with water before staining. Transgenic roots were selected based on DSred fluorescence using an Axiozoom (Zeiss). For staining, each root system was treated with 10% KOH for three days at room temperature, washed thrice with water, and stained with an ink solution (5% Schaeffer black ink, 95% acetic acid) for 10 min at 95°C. Stained roots were then soaked overnight in water and colonization quantified using the grid intersect method.^83^

## Supplementary Figure and Table Legends

**Figure S1: Unrooted phylogenetic tree of GPATs among 438 Viridiplantae species (log-likelihood: -848917.100).** Leaves are colored according to their lineage (orange: tracheophytes, green: bryophytes, blue: charophytes, pink: chlorophytes and cyan: prasinodermophytes). Genes from *Arabidopsis thaliana*, *Marchantia polymorpha*, *M. paleacea* and RAM2 from *Medicago truncatula* are indicated.

**Figure S2. Permeability to toluidine blue of plantlets from *Arabidopsis* mutant lines *gpat4,8* expressing different p-GPATs genes**. Plants (three independent lines) were cultivated either in vitro (left panel) or in soil in a greenhouse (right panel) before been assayed. At, *Arabidopsis thaliana*; Mpo, *Marchantia polymorpha*; Mpa, *Marchantia paleacea*.

**Figure S3. Permeability to toluidine blue of flowers from *Arabidopsis* mutant lines *gpat6* expressing different p-GPATs** genes. Three independent lines were assayed for each construct. At, *Arabidopsis thaliana*; Mpo, *Marchantia polymorpha*.

**Figure S4. Marchantia polymorpha RAM2 genes are expressed where cutin is deposited.** (A) Bright-field microcopy images showing*RAM2* promotors activities visualized by blue GUS staining in transgenic *M. polymorpha* lines and (B) the corresponding cross-sections at three weeks of age. GUS activities driven by the promoters of the *MpoRAM2* genes were observed in five independent GUS lines. Scale bar = 2mm for whole thalli and 600µm for cross-sections

**Figure S5. Mutation in the *Marchantia paleacea* the *ram2b*, *ram2c* and *ram2bram2c* mutants.**

**Figure S6. Growth phenotypes of the *Mparam2b* and *Mparam2c* mutants.** *Marchantia paleacea* gemmae of 3 independent mutant alleles were grown on 1/2 Gamborg media. While some lines showed growth delay, it was not consistant across alleles of the same genes.

**Figure S7. Scanning electron micrographs of the dorsal face of wild-type, *ram2b*, *ram2c* and *ram2b/ram2c* lines of *Marchantia paleacea***. Three independent lines were observed for each.

**Figure S8. Epidermis of wild-type, *ram2b*, *ram2c* and *ram2b/ram2c* lines of *Marchantia paleacea* as seen by transmission electron microscopy.** A. Three independent lines were observed for each. (Inset) Magnified image showing one of the rare epidermis cells with a cuticle. B. Cuticle width for the same lines (n=36 ± SD).

**Figure S9. Analysis of the *M. polymorpha SBG_pro_:GUS* construct in *M. paleacea*.** Promoter activity of the *M. polymorpha SBG* gene in *M. paleacea* plants monitored using Mpo*SBGpro:GUS* at 7 weeks post inoculation. From left to right, mock inoculated (brightfield), *R.irregularis*-inoculated (brightfield), *R.irregularis*-inoculated (WGA-Alexafluor 488), *R.irregularis*-inoculated (merged), *R.irregularis*-inoculated (merged magnified). Scale bar: 250 μm and 100 µm for magnified images. Three independent lines are shown.

**Figure S10. Streptophyte algae lack cutin.** Cutin monitoring by confocal microscopy in cross sections of *Marchantia polymorpha* thalli (Tak-1 accession) and streptophyte algae. Calcofluor and Nile red staining mark the primary cell wall and lipid contents, respectively. In *M. polymorpha*, lipid staining highlights the cutin layer (CL) coating the inner surface of the air chambers (AC) and the air pores (AP), whereas in streptophyte algae, the staining reveals intracellular lipid droplets. Scale bar = 20µm.

**Figure S11. Mycorrhizal phenotyping of the *Marchantia paleacea* single *ram2* mutants.** A. Exclusion zone, and colonizaion rate (NM= non-mycorrhized thalli; Myc= mycorrhized thalli. B. Ink-stained sections of the wild-type transformed with an Empty-Vector (EV) control, and the various *ram2b* and *ram2c* mutants.

**Table S1. List of species used for phylogeny + genome version + ref genomes.**

**Table S2. List of domain annotation per sequence.**

**Table S3. Quantification of cutin monomers in Marchantia wt and mutants.**

**Table S4. Quantification of cutin monomers in streptophyte algae and Embryophytes.**

**Table S5. List of primers used, and genetic constructs generated in this work.**

