## Supplementary material for "A single genetic innovation at the origin of plant terrestrialization": Figure S1 to S11

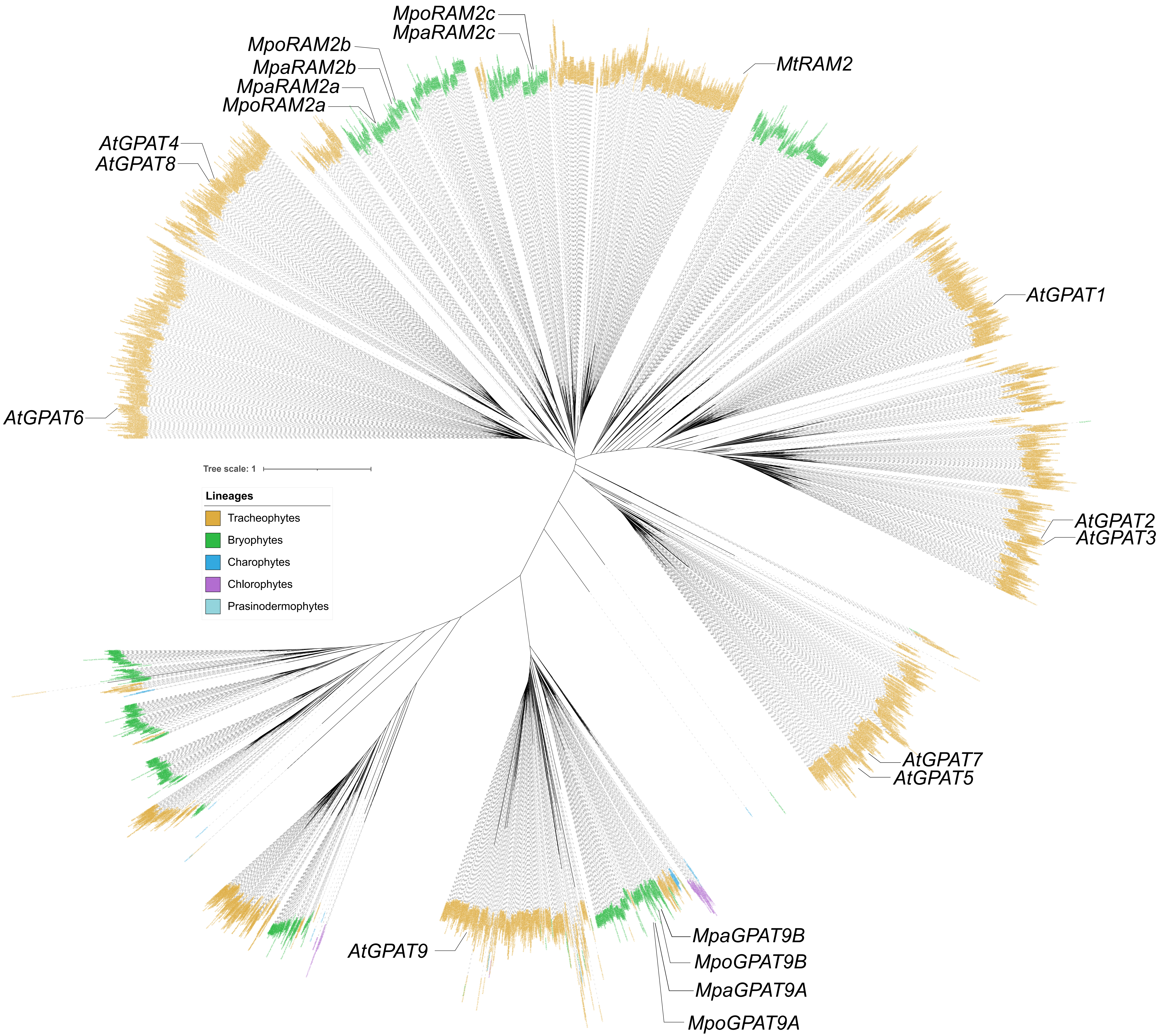


**Figure S1**: High-resolution full phylogeny

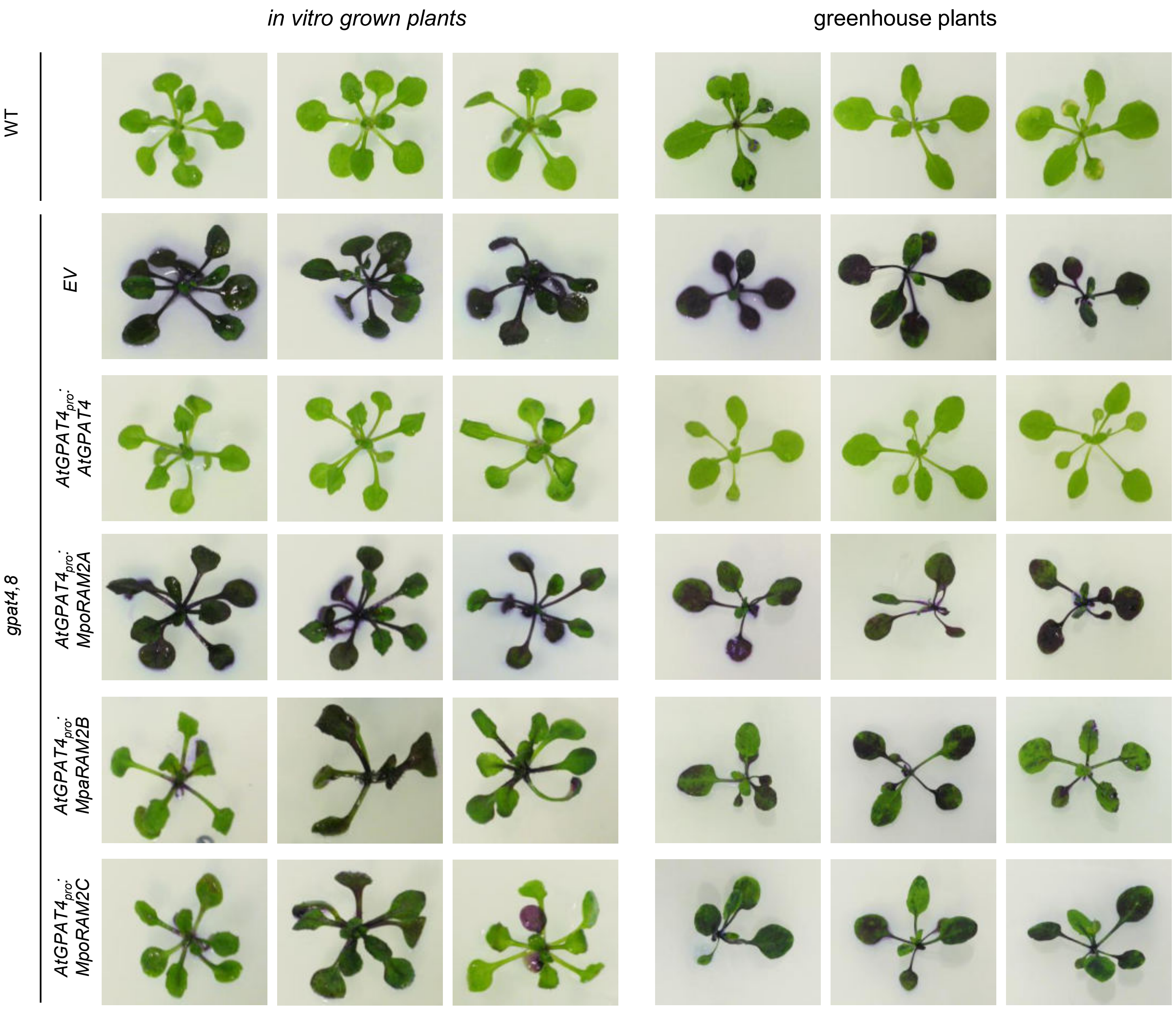


**Figure S2. Permeability to toluidine blue of plantlets from *Arabidopsis* mutant lines *gpat4,8* expressing different GPAT-like genes.**
Plants (three independent lines) were cultivated either in vitro (left panel) or in soil in a greenhouse (right panel) before been assayed. At, *Arabidopsis thaliana*; Mpo, *Marchantia polymorpha*; Mpa, *Marchantia paleacea*.

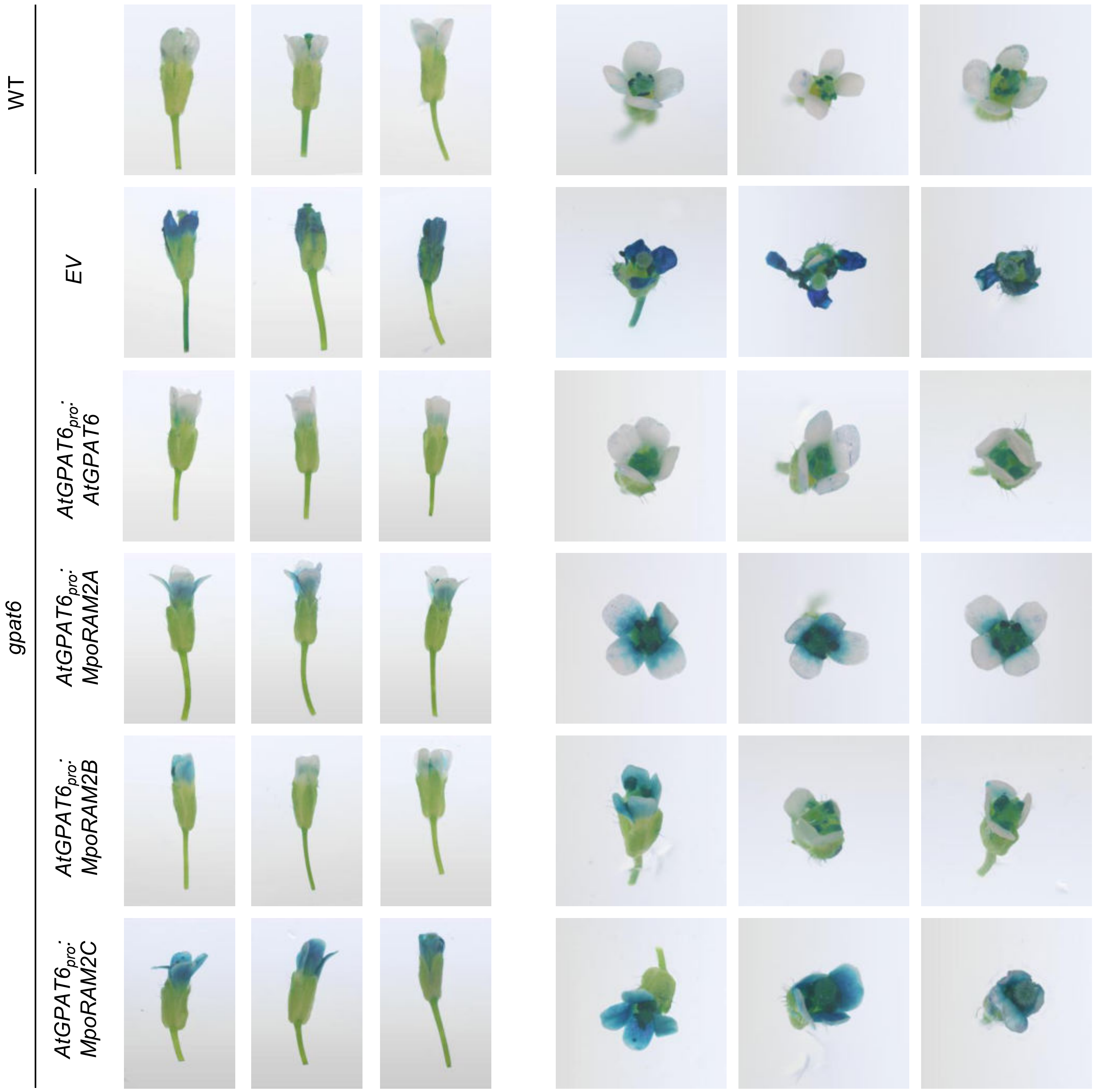


**Figure S3. Permeability to toluidine blue of flowers from *Arabidopsis* mutant lines *gpat6* expressing different GPAT-like genes.**
Three independent lines were assayed for each construct. At, *Arabidopsis thaliana*; Mpo, *Marchantia polymorpha*.

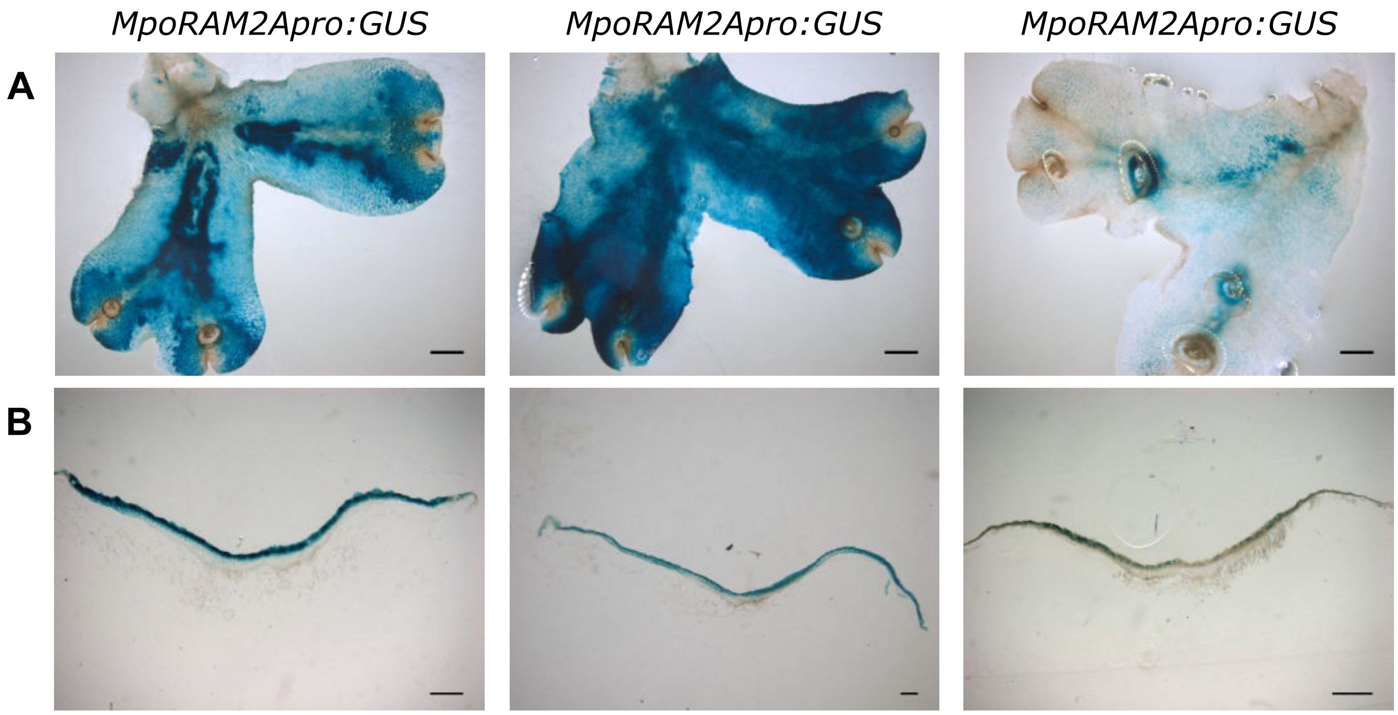


**Figure S4. *Marchantia polymorpha RAM2* genes are expressed where cutin is deposited.** (A) Bright-field microcopy images showing *RAM2* promotors activities visualized by blue GUS staining in transgenic *M. polymorpha* lines and (B) the corresponding cross-sections at three weeks of age. GUS activities driven by the promotors of the *MpoRAM2* genes were observed in five independent GUS lines. Scale bar = 2mm for whole thalli and 600µm for cross-sections.

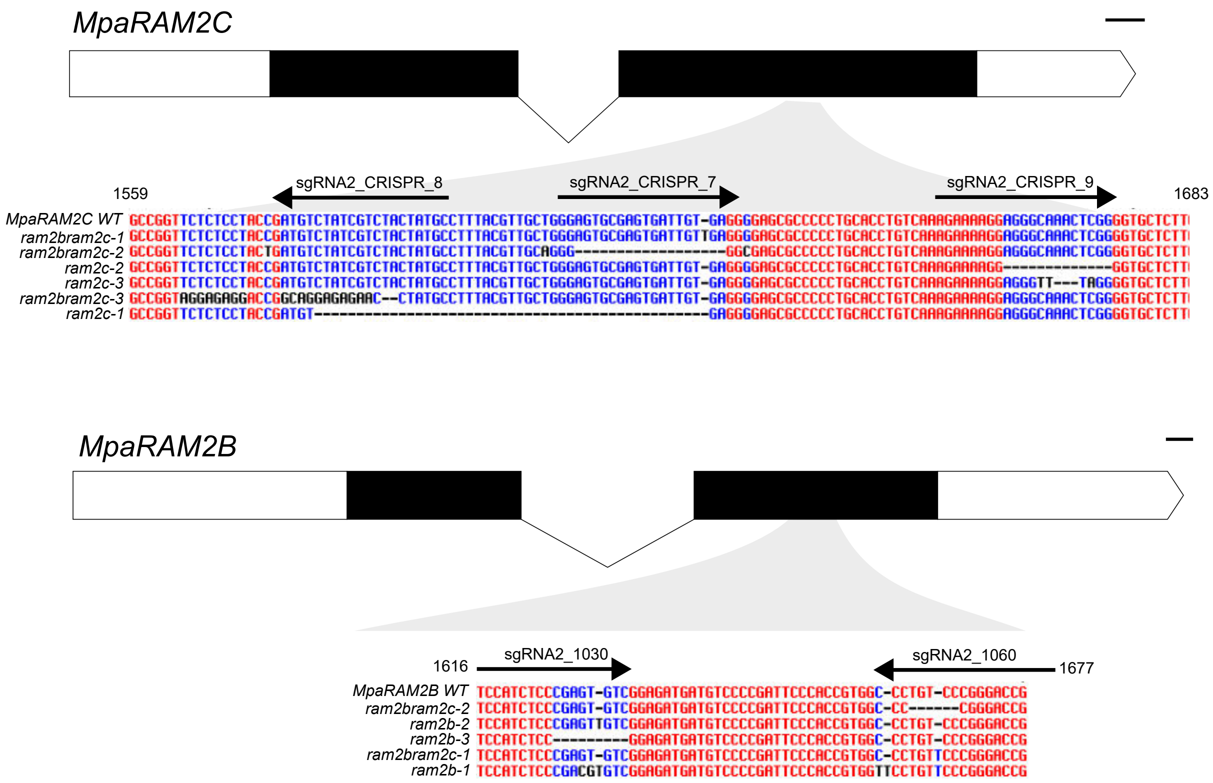


**Figure S5. Mutation in the *Marchantia paleacea ram2b*, *ram2c* and *ram2bram2c* mutants.**

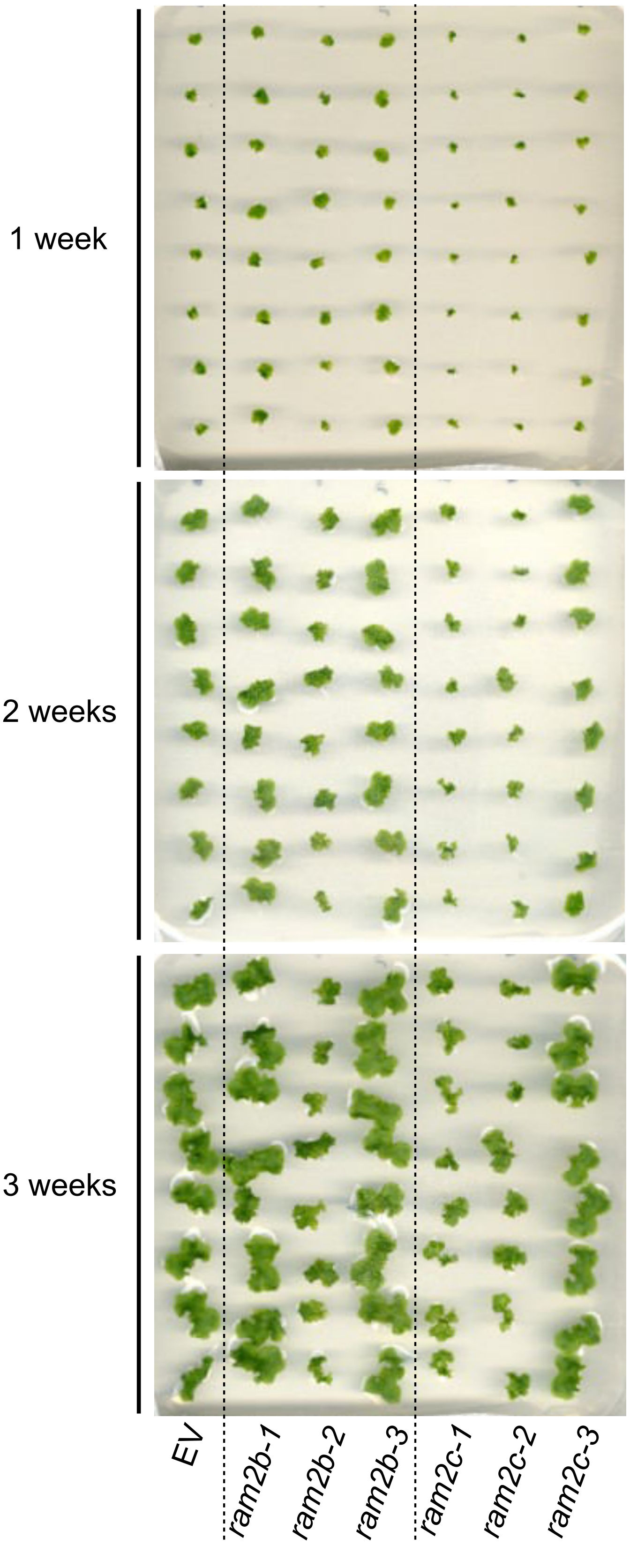


**Figure S6**: Growth phenotypes of the *Mparam2b* and *Mparam2c* mutants. *Marchantia paleacea* gemmae of 3 independent mutant alleles were grown on 1/2 Gamborg media. While some lines showed growth delay, it was not consistant across alleles of the same genes.

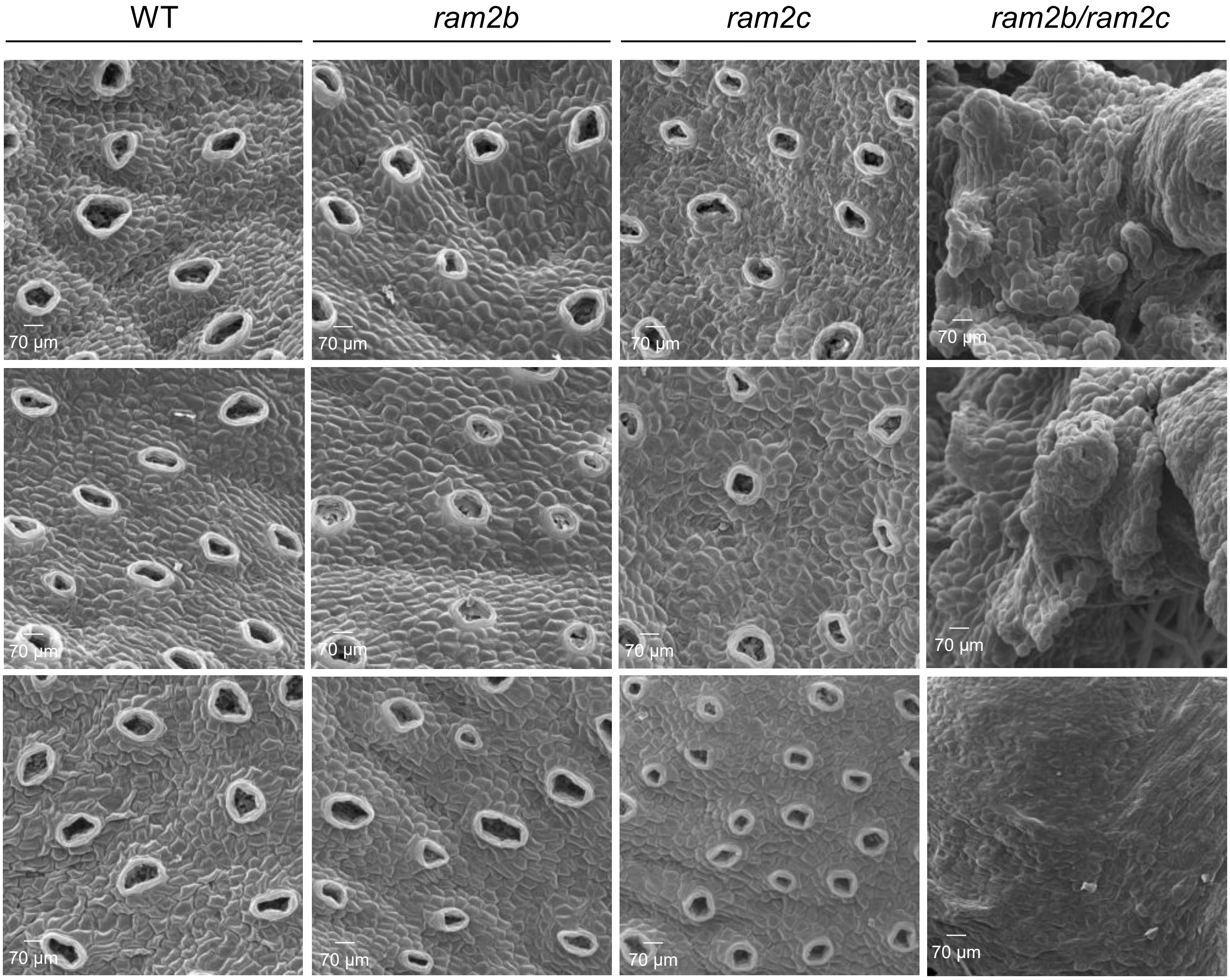


**Figure S7. Scanning electron micrographs of the dorsal face of wild-type, *ram2b*, *ram2c* and *ram2b/ram2c* lines of *Marchantia paleacea*.**
Three independent lines were observed for each.

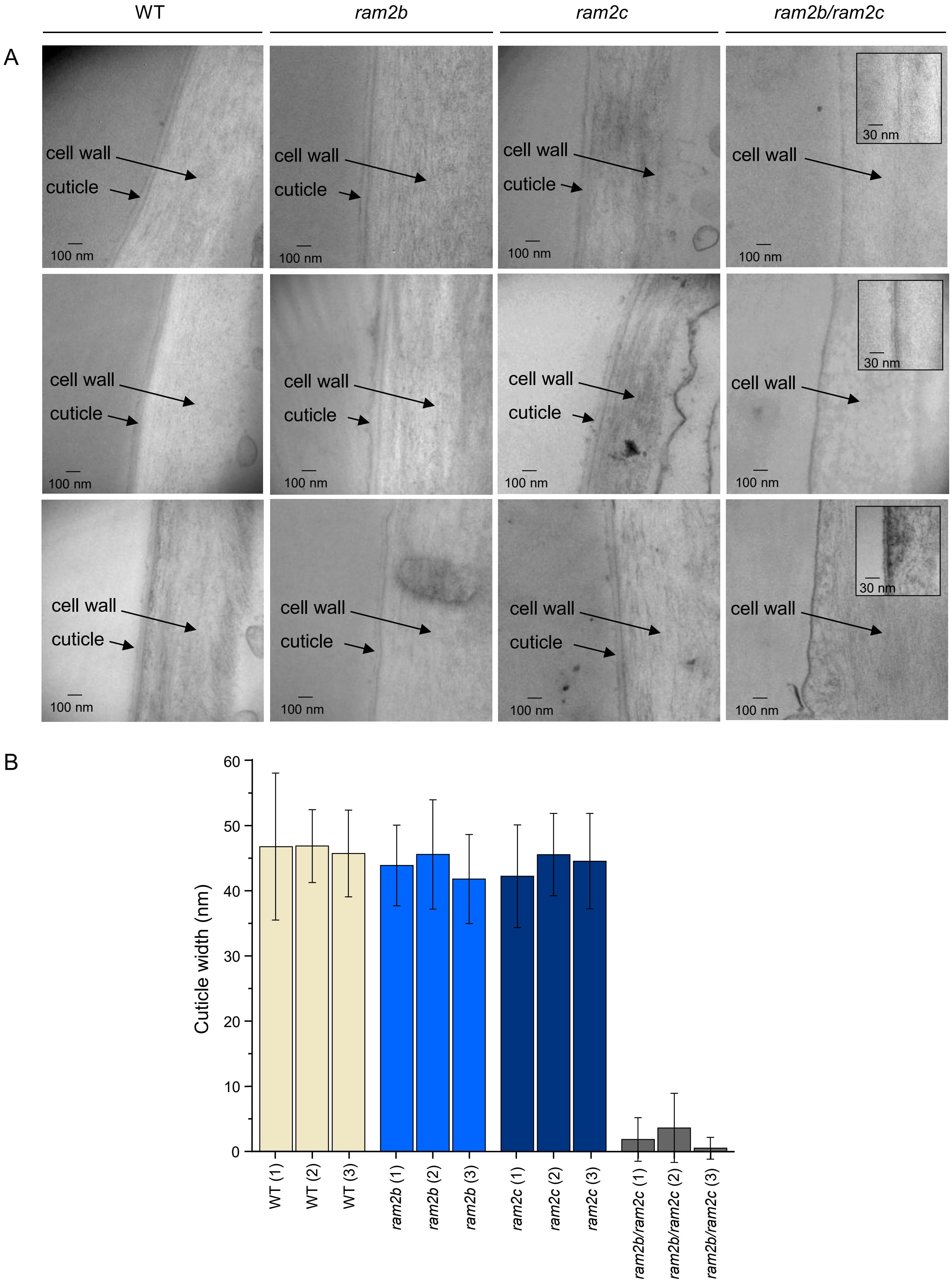


**Figure S8. Epidermis of wild-type, *ram2b*, *ram2c* and *ram2b/ram2c* lines of *Marchantia paleacea* as seen by transmission electron microscopy.**
A. Three independent lines were observed for each. (Inset) Magnified image showing one of the rare epidermis cells with a cuticle. B. Cuticle width for the same lines (n=36 ± SD).

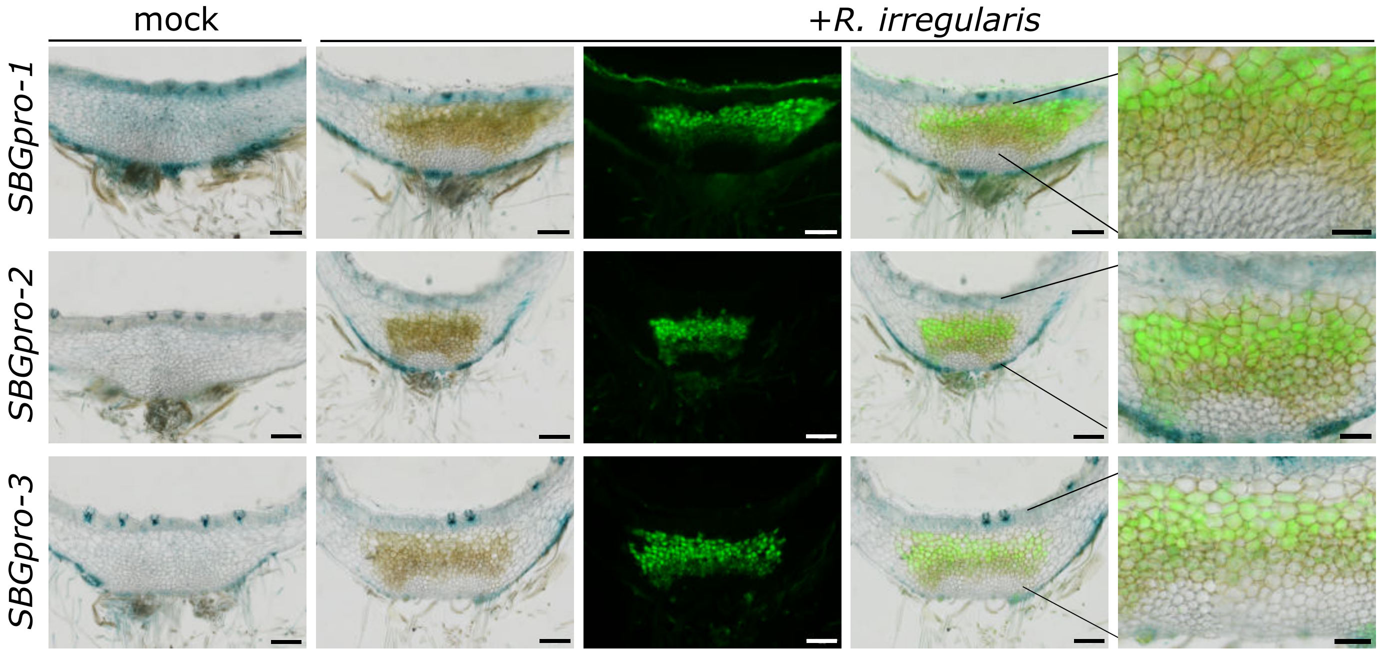


**Figure S9**: Analysis of the *M. polymorpha* SBGpro:GUS construct in *M. paleacea* Promoter activity of the *M. polymorpha* SBG gene in *M. paleacea* plants monitored using *MpoSBGpro:GUS* at 7 weeks post inoculation. From left to right, mock inoculated (brightfield), *R.irregularis*-inoculated (brightfield), *R.irregularis*-inoculated (WGA-Alexafluor 488), *R.irregularis-inoculated* (merged), *R.irregularis*-inoculated (merged magnified). Scale bar: 250 µm and 100 µm for magnified images. Three independent lines are shown.

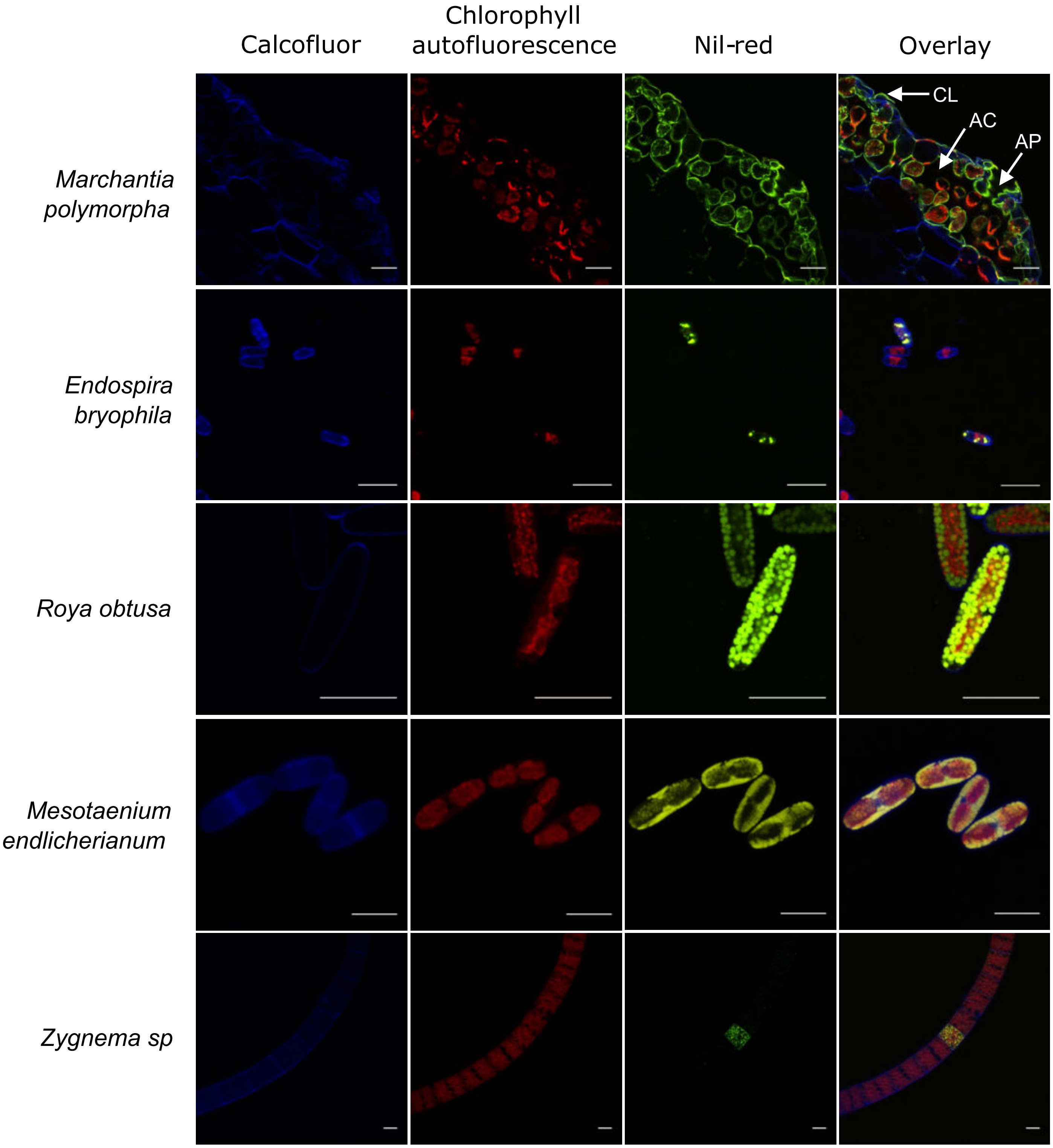


**Figure S10. Streptophyte algae lack cutin.** Cutin monitoring by confocal microscopy in cross sections of *Marchantia polymorpha* thalli (Tak-1 accession) and streptophyte algae. Calcofluor and Nil-red staining mark the primary cell wall and lipid contents, respectively. In *M. polymorpha*, lipid staining highlights the cutin layer (CL) coating the inner surface of the air chambers (AC) and the air pores (AP), whereas in streptophyte algae, the staining reveals intracellular lipid droplets. Scale bar = 20μm.

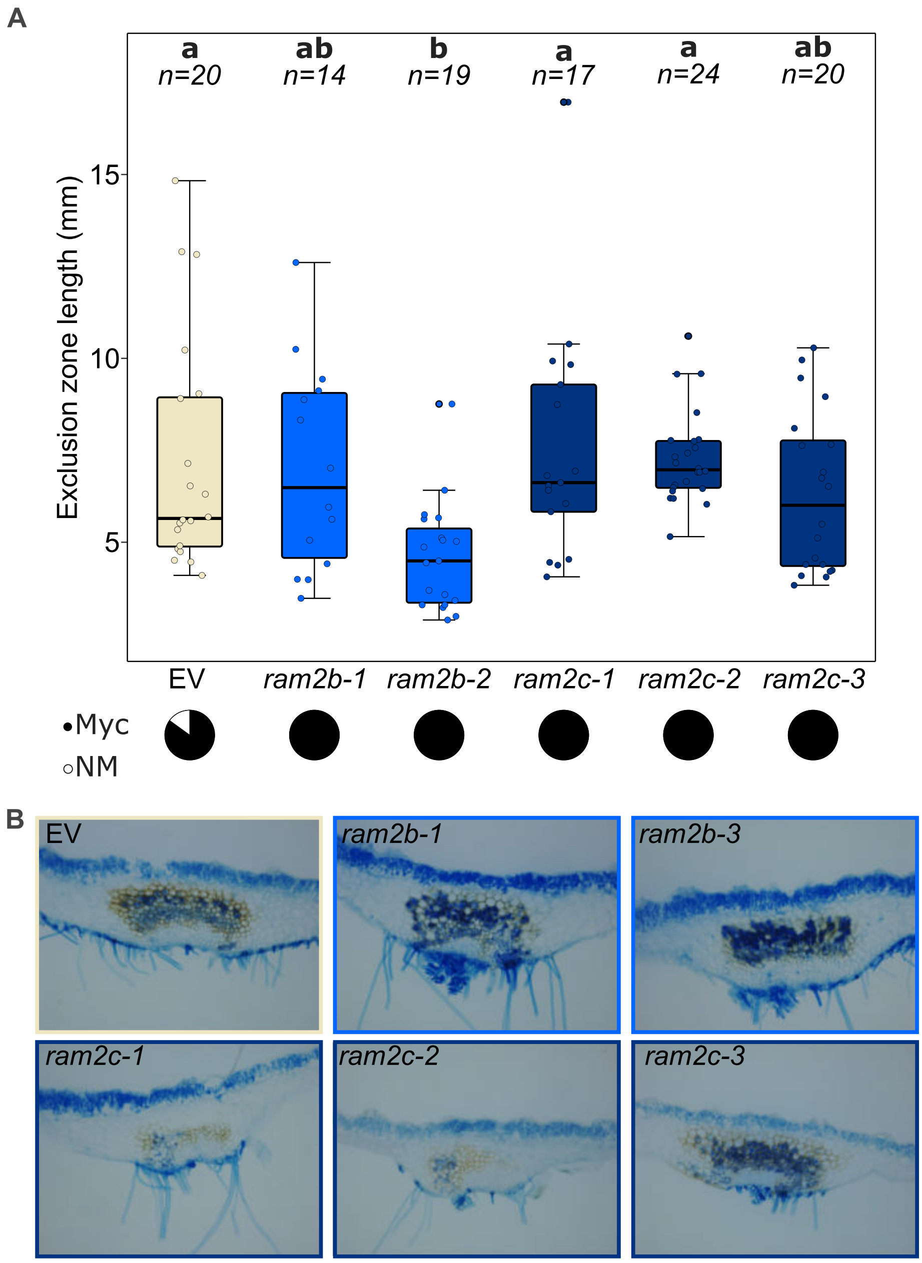


**Figure S11. Mycorrhizal phenotyping of the Marchantia paleacea single ram2 mutants.**
A. Exclusion zone, and colonizaion rate (NM= non-mycorrhized thalli; Myc= mycorrhized thalli
B. Ink-stained sections of the wild-type transformed with an Empty-Vector (EV) control, and the various *ram2b* and *ram2c* mutants.
